# Modifying meiotic recombination by targeting chromatin modifiers to crossover hotspots in *Arabidopsis*

**DOI:** 10.1101/2025.06.23.661042

**Authors:** Wojciech Dziegielewski, Maja Szymanska-Lejman, Anna Wilhelm, Karolina Hus, Piotr A. Ziolkowski

**Author notes:** These authors contributed equally to this work (Wojciech Dziegielewski, Maja Szymanska-Lejman).

## Abstract

The impact of specific chromatin modifications on meiotic crossover frequency is usually inferred from correlative studies, leaving open the question of causality. To address this, we used a dCas9-based system to target histone H3 methylation modifiers to defined genomic loci. Targeting methyltransferases responsible for H3K9me3 and H3K27me3 had little effect on recombination at selected hotspots, whereas targeting H3K4me3- and H3K36me3-associated enzymes often silenced the endogenous genes. The strongest effect was observed with the demethylase JMJ14, which reduced local H3K4me3 levels and decreased crossover frequency within the targeted interval. This was accompanied by reduced transcription of a long non-coding RNA (lncRNA) located at the hotspot and altered crossover topology. Suppressed recombination was also seen at neighbouring, untargeted hotspots. Conversely, directing the transcriptional activator VP64 to the same region elevated lncRNA expression, increased crossover frequency, and raised H3K4me3 levels. Our results reveal a causal relationship between H3K4me3, transcription, and local crossover activity, demonstrating that H3K4me3 levels are tightly associated with both transcriptional output and recombination frequency at specific genomic sites.

## Introduction

Meiotic crossovers represent the most powerful tool available to plant breeders, enabling the combination of beneficial genetic variation to develop cultivars with improved agronomic traits. Typically, however, breeders are interested in transferring only specific loci associated with desirable traits between parental genotypes. This is particularly important for the introgression of disease resistance genes, which are often tightly linked to unfavourable alleles. Unfortunately, no current technology allows for the targeted induction of recombination at specific genomic locations. As a result, breeding programs rely on screening large plant populations, making the process both time-consuming and resource-intensive.

Meiotic recombination is initiated through the formation of programmed DNA double-strand breaks (DSBs), catalyzed by the evolutionarily conserved transesterase SPO11, in conjunction with the topoisomerase MTOPVIB (Bergerat *et al*, 1997; Keeney *et al*, 1997; Grelon *et al*, 2001; Hartung *et al*, 2007; Fu *et al*, 2016; Vrielynck *et al*, 2016). In most eukaryotes, including plants, only a small fraction of meiotic DSBs is resolved into crossovers, while the majority are repaired through alternative non-crossover pathways. For example, in *Arabidopsis thaliana*, out of approximately 200 DSBs formed during a single meiotic division, no more than 10 are repaired as crossovers (Ferdous *et al*, 2012; Wijnker *et al*, 2013). It is possible that this large ‘excess’ of DSBs underlies the failure of DSB induction-based strategies to enhance crossover formation at specific loci in plants, unlike in yeast (Yelina *et al*, 2022).

The distribution of COs along chromosomes is non-uniform but instead occurs preferentially at specific regions known as recombination hotspots, where the crossover frequency significantly exceeds the genome-wide average (Wu & Lichten, 1995; Myers *et al*, 2005; Pan *et al*, 2011; Choi *et al*, 2013). In the majority of mammals such as mice and humans, the location of most DSB hotspots is determined by PRDM9, a histone methyltransferase containing a zinc finger DNA-binding domain that recognizes specific sequence motifs (Baudat *et al*, 2010; Yamada *et al*, 2017; Diagouraga *et al*, 2018). PRDM9 directs the deposition of histone modifications, including trimethylation of histone H3 at lysines 4 (H3K4me3) and 36 (H3K36me3), which in turn facilitates the recruitment of the DSB machinery (Baudat *et al*, 2010; Diagouraga *et al*, 2018). Plants lack PRDM9 and therefore employ alternative mechanisms to define recombination hotspots. One of the factors stimulating DSB formation in plants is chromatin accessibility as the majority of recombination hotspots are observed in low nucleosome density regions (LNDs), which corresponds to gene promoters and 3’ ends of the genes (Choi *et al*, 2013, 2018; He *et al*, 2017; Kianian *et al*, 2018). This coincides with deposition of H2A.Z histone variant at transcription start sites (TSS) and transcription termination sites (TTS). Furthermore, several studies have implicated specific histone marks in either promoting or repressing meiotic recombination. For instance, euchromatic features such as histone H3 lysine 4 dimethylation and trimethylation (H3K4me2, H3K4me3) as well as H3K9 acetylation have been associated with crossover hotspots in multiple species, while heterochromatic marks such as H3K9me2, H3K27me3 or H2A.W histone variant are generally enriched in recombination-poor regions (Borde *et al*, 2009; Buard *et al*, 2009; Yamada *et al*, 2013; Underwood *et al*, 2018; Son *et al*, 2025). However, the causal relationship between these chromatin states and recombination outcomes remains unclear, particularly in plants, where direct experimental evidence is limited. Consequently, understanding whether local manipulation of chromatin marks can modulate crossover activity is a critical step toward enabling targeted recombination in plant genomes.

In this study, we employed a CRISPR–dCas9 platform in *A. thaliana* to test influence of specific chromatin modifiers on meiotic recombination frequency in hotspots within a defined pericentromeric interval. By leveraging dCas9-mediated recruitment of methyltransferases affecting H3K4, H3K36, H3K9, and H3K27 to fluorescently tagged crossover hotspots, we establish a versatile and quantitative assay for probing recombination outcomes. Our findings highlight a critical role for H3K4me3 in promoting crossover activity, as demonstrated by the suppressive effect of the histone demethylase JUMONJI 14 (JMJ14). In contrast, enhancing transcription through VP64 activation significantly elevated crossover rates, underscoring a tight link between transcriptional activity and hotspot function.

## RESULTS

### A system for testing the effects of chromatin modification on crossover formation

In plants, recombination hotspots are relatively numerous and are primarily located in the 5′ and 3′ UTRs of most transcriptionally active genes (Choi *et al*, 2012; Drouaud *et al*, 2013; Wijnker *et al*, 2013; He *et al*, 2017; Choi *et al*, 2018; Kianian *et al*, 2018). As a result, determining the direct impact of chromatin state on the recombination activity of hotspots requires a system capable of measuring crossover frequency within short chromosomal regions containing as few hotspots as possible. We recently developed a system based on extremely short chromosomal intervals (ESILs), which are defined by fluorescent reporters expressed in seeds. By scoring the segregation of these reporters during meiosis, it is possible to precisely measure crossover frequency within the interval (Fig. 1a). Since ESIL intervals span only a few dozen kilobases and contain just a few hotspots, this system is ideal for directly studying how local changes in chromatin state affect crossover frequency.

**Figure 1.**
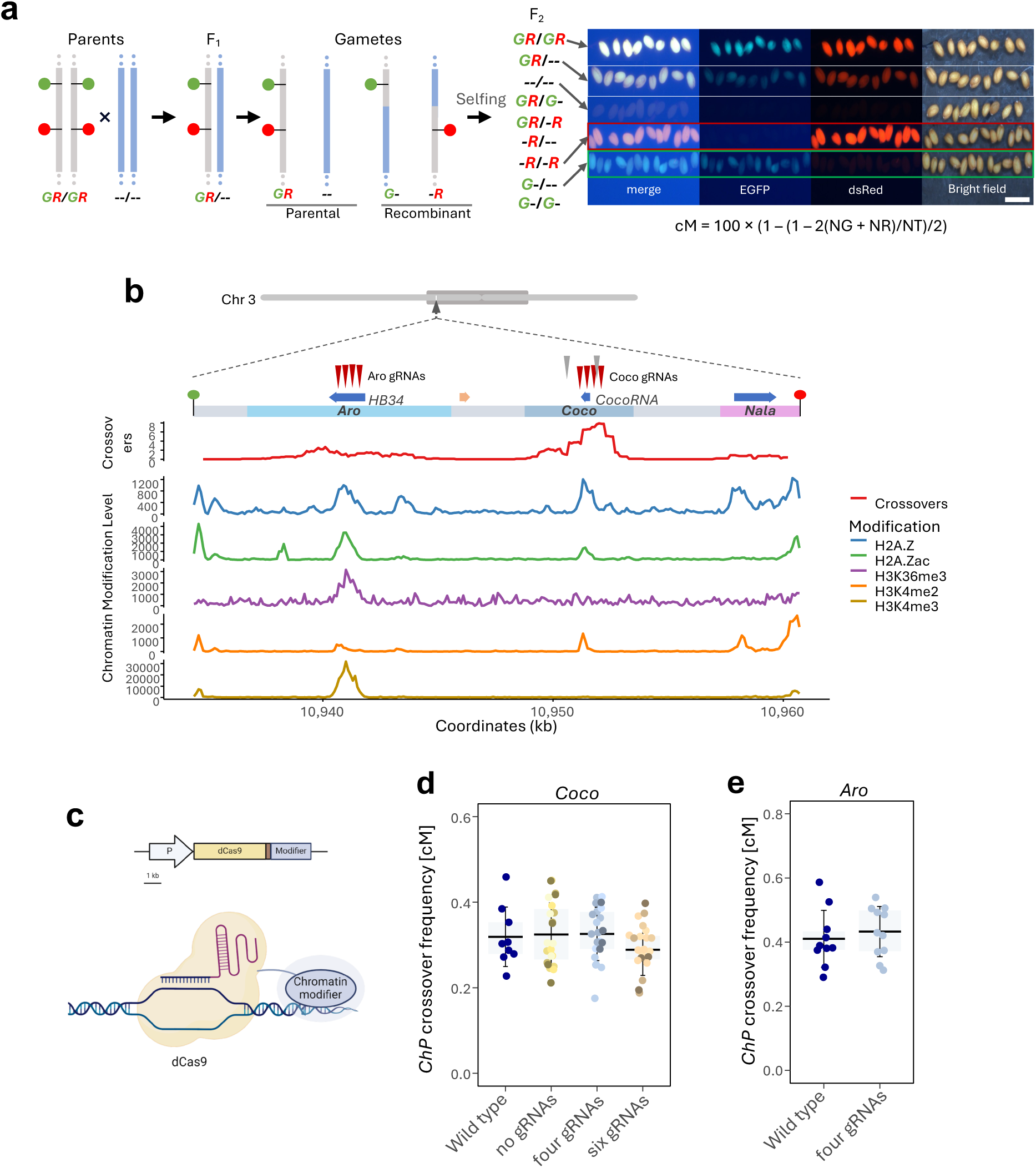
System for studying the impact of chromatin modifications on crossover frequency. **a.** Measurement of crossover frequency based on ESILs with segregating fluorescent reporters. The vertical gray and blue lines represent homologous chromosomes from Col (ESIL) and L*er*, respectively. Green and red markers indicate fluorescent reporters. All possible F_2_ genotypes are shown. On the right, representative images of seeds from the Col-*ChP* × L*er* cross are displayed. **b.** Chromosomal location, genes, and chromatin landscape of the *ChP* interval. The top panel shows the position of *ChP* on the chromosome, with the pericentromeric region indicated by a dark gray oval. A zoomed-in view highlights the interval, marking the *Aro*, *Coco*, and *Nala* crossover hotspots. Blue horizontal arrows indicate gene locations, while an orange arrow marks a pseudogene. Vertical maroon arrowheads indicate the approximate positions of the gRNAs used, while gray arrowheads mark gRNAs employed exclusively in the steric hindrance experiment (six-gRNA version). Crossover frequencies in the wild type are plotted as a moving average in a 1 kb window with a 100 bp step. Levels of specific chromatin modifications are displayed in 100 bp windows. Crossover data based on Szymanska-Lejman et al. (2023), H2A.Z and H2A.Zac levels from Bieluszewski et al. (2022), H3K36me3 from Liu et al. (2021), H3K4me2 from Yu et al. (2021), H3K4me3 from Potok et al. (2022). **c.** A schematic representation of the constructs used for dCas9-modifier expression and the mode of action of the resulting fusion protein. **d-e**. dCas9 without additional effectors does not affect crossover frequency in the *ChP* interval. **d.** Crossover frequency in crosses with transformants expressing dCas9 without gRNAs or with four/six gRNAs targeting the *Coco* hotspot. The Col-*ChP* × L*er* cross served as the wild-type control. In the boxplot, the centre line indicates the median, the upper and lower bounds represent the 75th and 25th percentiles, and error bars show the standard deviation. Each data point represents a crossover frequency measurement for a single cross. At least three independent transformants were used in each case. **e.** As in (**d**), but for crosses with lines expressing dCas9 with four gRNAs targeting the *Aro* hotspot.

To test the impact of local chromatin modifications on meiotic recombination, we selected the *Chili Pepper* (*ChP*) interval, which contains three distinct crossover hotspots (Fig. 1b). To locally modify chromatin within these hotspots, we employed a system based on a catalytically inactive form of the Cas9 nuclease (deadCas9, dCas9). In this system, dCas9 is fused to a selected effector via translational fusion. The recruitment of dCas9-effectors to specific genomic loci is guided by CRISPR guide RNAs (gRNAs), expressed under *U3* or *U6* promoters (Fig. 1c).

Since dCas9 itself is a large protein (156 kDa), we considered the possibility that targeting dCas9 to a hotspot might cause steric hindrance, limiting the access of recombination proteins and thus reducing crossover frequency. To test this, we directed dCas9 independently to two hotspots within the *ChP* interval, *Coco* and *Aro*, using either four or six gRNAs that were complement to the highest recombination peaks in both hotspots (Fig. 1b). The transformants generated in the Col-*ChP* genetic background were then crossed with wild-type plants (L*er*-0), and crossover frequency was measured for each. The results showed that recruitment of dCas9 without an effector to either *Coco* or *Aro*, had no effect on crossover formation at the *ChP* interval (Fig. 1d,e). Similar results were obtained when T_1_ lines carrying gRNAs targeting the *Coco* hotspot were crossed to Col (Appendix Fig. S1). Based on this, we concluded that recombination activity within hotspots is not altered due to steric hindrance caused by dCas9 targeting.

### A screen for H3 histone modification revealed JMJ14 histone demethylase as having the effect on hotspot recombination activity

To investigate whether local chromatin modifications influence hotspot activity, we generated a set of genetic constructs in which the catalytic domains or full coding sequences of chromatin effectors were fused to catalytically inactive Cas9 (dCas9). The specific fragments of each effector used for dCas9 fusions are presented in Appendix Fig. S2 and Fig. 2b. The schematic workflow of the experiment is showed in Appendix Fig. S3. Targeting to the recombination hotspots, located within the *ChP* interval, was achieved using a set of four guide RNAs (gRNAs) previously validated in the steric hindrance assay. As a control, we used constructs with functional dCas9-effectors but without gRNAs, thereby preventing recruitment to the target hotspots. Genetic constructs were introduced into Col-*ChP* plants and crossed with the non-fluorescent accession, L*er*, to assess crossover frequency by fluorescent seed segregation (Appendix Fig. S3).

**Figure 2.**
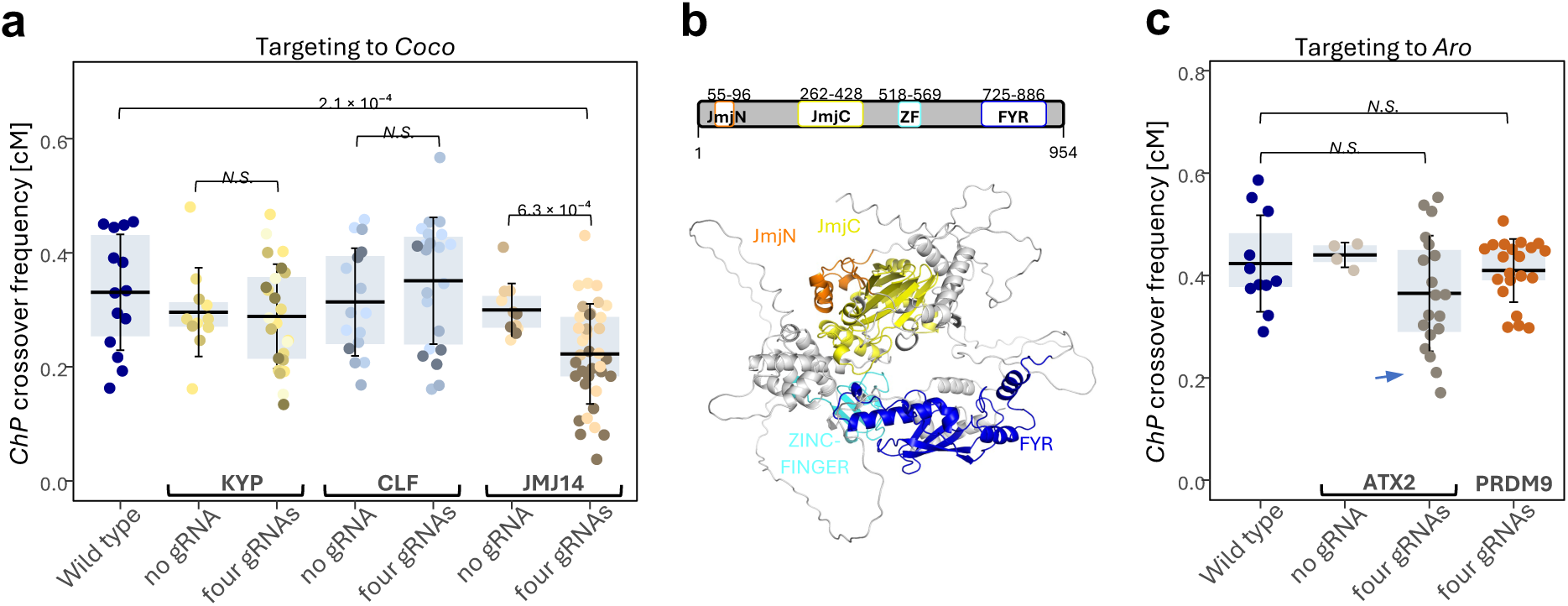
Screening for chromatin modifiers reveals the histone demethylase JMJ14 as a protein capable of locally reducing crossover frequency through CRISPR-dCas9 targeting. **a.** Crossover frequency in the *ChP* interval in crosses with transformants expressing dCas9 fused to the SET domains of histone methyltransferases KYP, CLF, or the histone demethylase JMJ14, targeted to the *Coco* hotspot. Controls included plants without gRNAs and wild-type Col-*ChP* × L*er*. In the boxplot, the center line indicates the median, the upper and lower bounds represent the 75th and 25th percentiles, and error bars show the standard deviation. Each data point represents crossover frequency for a single cross. At least three independent transformants were analyzed per construct. **b.** Domain architecture and predicted structure of the JMJ14 protein. The upper panel illustrates the domain organization of JMJ14, highlighting key functional regions: JmjN (Jumonji N-terminal domain), JmjC (Jumonji C-terminal domain), ZF (Zinc Finger domain), and FYR (Phenylalanine–Tyrosine–Rich domain). The lower panel shows a structural model of JMJ14 predicted by AlphaFold2. The JmjN and JmjC domains form the catalytic core responsible for histone demethylation. The ZF domain is implicated in DNA binding, while the FYR domain is likely involved in protein–protein interactions. **c.** As in (**a**), but for targeting the SET domain of histone methyltransferase ATX2 and the full-length human PRDM9 protein to the *Aro* hotspot. The blue arrow indicates crosses with potentially silenced *ATX2*.

Histone methylations H3K9me2/3 and H3K27me3 correlate with transcriptional gene repression and serve as classic markers of “closed chromatin” (reviewed in Lloyd & Lister, 2022). It is widely believed that closed chromatin limits the formation of meiotic DSBs by SPO11 as well as crossover formation (Pan *et al*, 2011; Choi *et al*, 2018; Underwood *et al*, 2018). Therefore, we first tested whether reducing H3K9me2/3 or H3K27me3 levels within the recombinationally hyperactive hotspot *Coco* could locally decrease crossover frequency. We fused the catalytic domains of the KRYPTONITE/SUPPRESSOR OF VARIEGATION HOMOLOG 4 (KYP/SUVH4) methyltransferase, responsible for H3K9me2 (Jackson *et al*, 2002; Malagnac *et al*, 2002), and CURLY LEAF (CLF), responsible for H3K27me3 (Goodrich *et al*, 1997; Schubert *et al*, 2006) to dCas9 (Appendix Fig. S2a,b), however, we did not observe a reduction in crossover frequency within the *ChP* interval after targeting either dCas9-KYP or dCas9-CLF to the *Coco* hotspot (Fig. 2a).

In many eukaryotes, H3K4 di/trimethylation (H3K4me2/3) positively correlates with crossover frequency and is enriched at recombination hotspots (Borde *et al*, 2009; Choi *et al*, 2013; Buard *et al*, 2009; Acquaviva *et al*, 2013). Therefore, we aimed to determine whether removing these modifications from nucleosomes within the *Coco* hotspot would reduce recombination activity (see Fig. 1b for H3K4me2/3 profile over *Coco* hotspot). We used JUMONJI 14 (JMJ14), an enzyme capable of demethylating H3K4me3 and, to a lesser extent, H3K4me2 in vivo (Lu *et al*, 2010; Yang *et al*, 2010). Targeting dCas9-JMJ14 to *Coco* led to a significant decrease in crossover frequency within the *ChP* interval, from 0.33 cM in the wild type to 0.22 cM in lines with JMJ14 targeting (Fig. 2a,b; *P* = 2.1 × 10⁻⁴, Welch test). Importantly, this effect was not observed in control lines lacking gRNAs, where JMJ14 was not recruited to *Coco* (Fig. 2a).

Next, we aimed to determine whether targeting the SET domains of H3K4me2/3 methyltransferases could increase crossover frequency in *ChP*. Since *Coco* exhibits very high native recombination activity, we instead targeted a weaker hotspot within *ChP*, *Aro* (Fig. 1b). We generated dCas9 constructs fused with the catalytic domains of various methyltransferases: Arabidopsis TRITHORAX 2 (ATX2), responsible for H3K4me2/3 methylation (Saleh *et al*, 2008); SET DOMAIN GROUP 4 (SDG4), which methylates H3K4me2/3 and H3K36me3 (Schubert *et al*, 2006); and SET DOMAIN GROUP 8 (SDG8), which trimethylates H3 at K4 and K36 residues (Zhao *et al*, 2005) (Appendix Fig. S2c-e). Additionally, we created dCas9 fusion constructs with the human PR/SET DOMAIN 9 (PRDM9), a meiosis-specific H3K4me2/3 and H3K36me3 methyltransferase (Hayashi *et al*, 2005; Smagulova *et al*, 2011; Grey *et al*, 2011) (Appendix Fig. S2f).

For ATX2 and PRDM9, we observed no changes in *ChP* crossover frequency (Fig. 2c). However, plants expressing dCas9-SDG4 showed a reduction in *ChP* crossovers, both in lines targeting *Aro* and in control lines without gRNAs (Appendix Fig. S4a). This suggested SDG4 silencing in dCas9-SDG4 lines that resulted in an overall decrease in methyltransferase activity. Indeed, RT-qPCR analysis confirmed that *SDG4* expression in these lines was reduced to 0.2–0.5 of wild-type levels (Appendix Fig. S4b,c). Moreover, some degree of silencing was also observed in dCas9-ATX2 lines (Fig. 2c, Appendix Fig. S5). In dCas9-SDG8 plants, both transformants with and without gRNAs were nearly completely sterile, suggesting even stronger silencing of SDG8 expression (Appendix Fig. S4d). Together, these results indicate that the main challenge with increasing H3K4 and H3K36 methylation is the significant suppression of endogenous methyltransferase expression. Nevertheless, the lack of effect on crossover frequency in PRDM9-expressing lines suggests that, at least at this hotspot, targeting H3K4/K36 methyltransferase alone may not be sufficient to influence recombination activity.

### Targeted JMJ14 reduces local H3K4me3 levels, leading to both reduced crossover activity and transcriptional repression

To further investigate how JMJ14 reduces recombination activity in the *ChP* interval, we examined dCas9-JMJ14 enrichment at the *Coco* and *Aro* hotspots (Fig. 3a). We selected three independent T_1_ lines, each crossed with L*er* that showed varying degrees of *ChP* crossover reduction. Additionally, we included a line lacking gRNAs as a control. ChIP-qPCR analysis, performed using an antibody against HA tag that is fused to the C-terminus of dCas9, confirmed dCas9 enrichment at the *Coco* hotspot in gRNA-targeted lines, relative to wild-type plants. No enrichment was detected in the line without gRNAs (Fig. 3b). Importantly, dCas9 was also absent at the *Aro* hotspot, which was not targeted by gRNAs (Fig. 3b). These results confirm the efficiency and specificity of dCas9-JMJ14 binding to chromatin regions defined by gRNA sequences.

**Figure 3.**
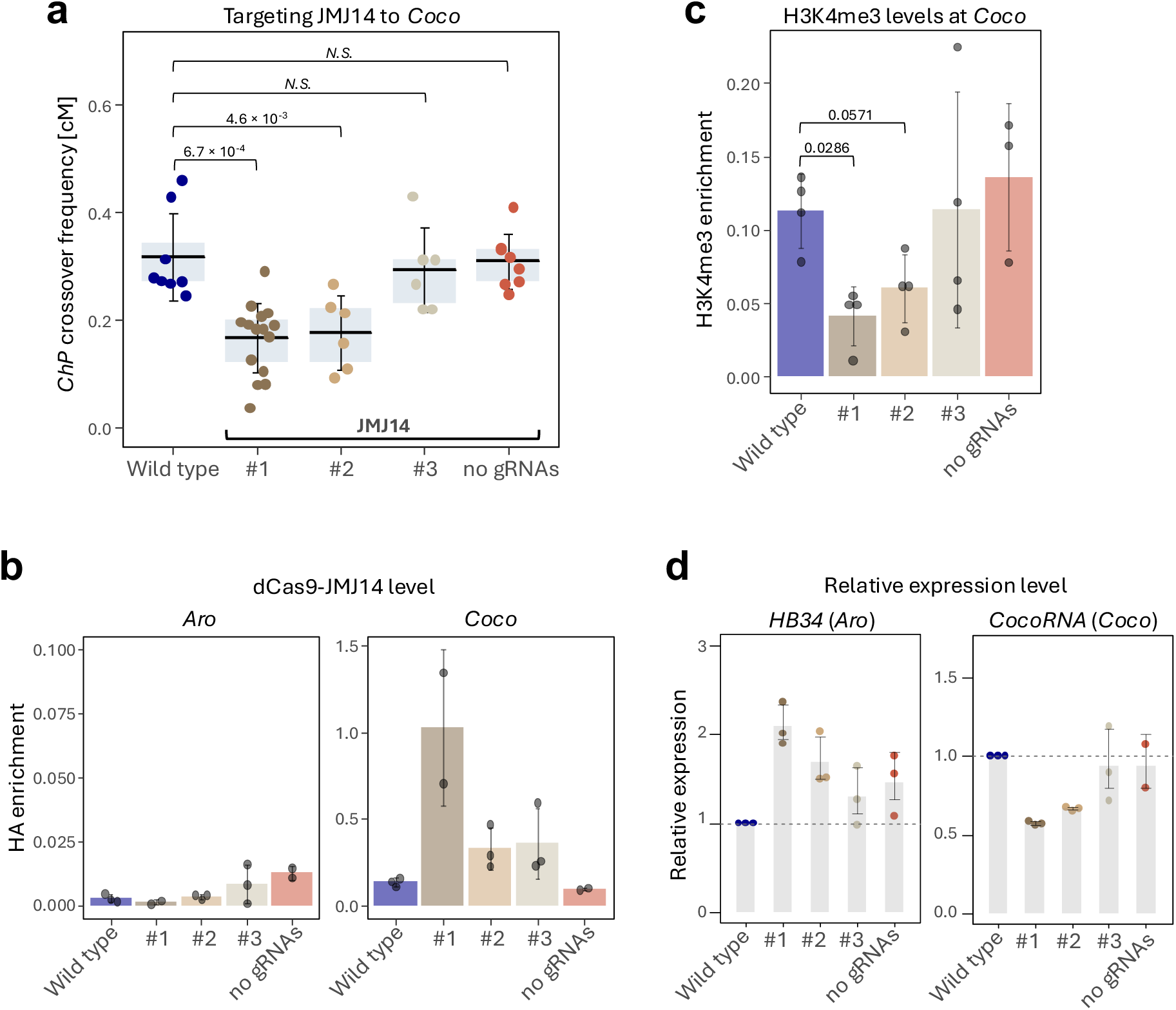
Targeted JMJ14 locally reduces crossover frequency and gene expression, associated with decreased H3K4me3 levels. **a.** Crossover frequency in the *ChP* interval for crosses with three transgenic lines carrying dCas9-JMJ14 targeted to the *Coco* hotspot (#1-3). Controls include wild-type Col-*ChP* × L*er* and crosses with lines carrying dCas9-JMJ14 but without gRNAs. In the boxplot, the center line indicates the median, the upper and lower bounds represent the 75th and 25th percentiles, and error bars show the standard deviation. Each data point represents crossover frequency for a single cross. Welch’s test was used to determine statistical significance. **b.** dCas9-JMJ14 enrichment at the *Aro* and *Coco* hotspots in the crosses used in (**a**), measured by ChIP-qPCR. Bars represent mean values of three biological replicates (dots), with error bars indicating standard deviation. **c.** Changes in H3K4me3 enrichment at the *Coco* hotspot in the crosses used in (**a**), measured by ChIP-qPCR. Bars represent mean values of three biological replicates (dots), with error bars indicating standard deviation. Statistical significance was assessed using Mann-Whitney U test. **d.** Relative expression of the *HB34* gene located within the *Aro* hotspot and the *CocoRNA* gene within the *Coco* hotspot in the crosses used in (**a**), measured by RT-qPCR. Bars represent mean values of three biological replicates (dots), with error bars indicating standard deviation.

Next, we examined H3K4me3 enrichment in these lines at the targeted hotspot (Fig. 3c). As expected, H3K4me3 levels were significantly reduced to 0.25-0.5 of wild-type levels in two lines exhibiting a decrease in *ChP* crossover frequency (*P* = 0.0266; Mann Whitney U test). However, in the third line, where dCas9-JMJ14 was detected but no significant change in crossover frequency was observed, H3K4me3 levels remained unchanged (Fig. 3a-c). In contrast, in the line lacking gRNAs, H3K4me3 methylation was higher than in the wild type. These findings clearly support the conclusion that the reduction in H3K4 trimethylation directly impacts hotspot recombination activity.

Similar to most crossover hotspots in Arabidopsis, the *Aro* and *Coco* hotspots overlap with actively transcribed genes: *HB34* (AT3G28920) and AT3G05605, respectively (Fig. 1b). AT3G05605 encodes an uncharacterized long noncoding RNA (lncRNA), which we named *CocoRNA*. Since decreased H3K4me3 levels often correlate with transcriptional repression, we investigated whether *CocoRNA* expression was reduced in dCas9-JMJ14 lines using RT-qPCR. Indeed, *CocoRNA* levels were significantly lower than in wild-type plants, correlating with the reduction in *ChP* crossover frequency (Fig. 3d). However, no changes in *CocoRNA* expression were observed in the line that did not show a crossover frequency reduction or in the line without gRNAs (Fig. 3d). Interestingly, *HB34* expression showed the opposite trend, increasing up to 2.1-fold in lines with reduced crossover frequency compared to wild-type plants, suggesting JMJ14-dependent coregulation of these two neighbouring genes (Fig. 3d). Together, these results demonstrate that histone demethylase targeting altered the chromatin state within the *Coco* hotspot, leading not only to a decrease in its recombination activity but also to transcriptional repression of the hotspot-associated gene.

### JMJ14 targeting alters crossover distribution within and between the three hotspots in the *ChP* interval

To precisely examine changes in the distribution of crossover events within the *ChP* interval in plants with targeted dCas9-JMJ14, we employed our recently developed seed-typing technique (Szymanska-Lejman *et al*, 2023). Seed-typing relies on deep sequencing of recombinant seeds selected from crosses with ESILs. A cross between a Col-*ChP* line carrying dCas9-targeted JMJ14 and a L*er* background generated a Col/L*er* hybrid, which, upon self-pollination, produced seeds segregating for eGFP and dsRed reporters. Recombinants were manually selected, and the crossover-containing interval was amplified using high-fidelity long-range PCR with overlapping amplicons. The resulting PCR products were used to construct Illumina-compatible libraries for each recombinant, followed by sequencing at ∼1500× coverage. Crossover sites were identified based on SNPs distinguishing Col from L*er*. By combining crossover positions from multiple recombinants, we achieved high-resolution mapping of crossover distribution (Fig. 4a).

**Figure 4.**
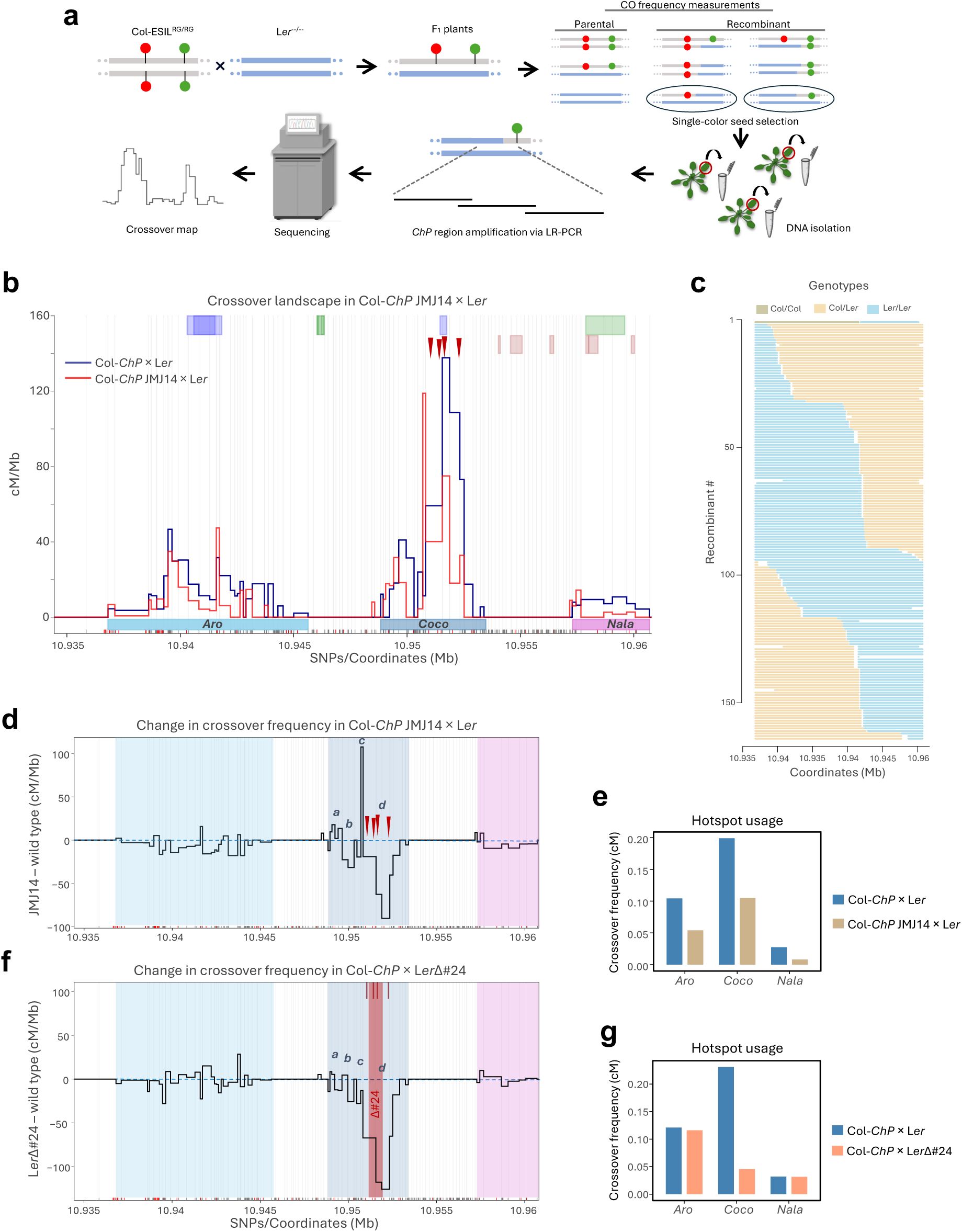
JMJ14 suppresses crossovers at the manipulated hotspot, alters its topology, and impacts adjacent recombination sites. **a.** Schematic overview of seed-based crossover mapping. A cross between ESIL and L*er* generates F_1_ individuals, in which crossover frequency within a defined interval marked by fluorescent reporters is measured by analyzing the segregation of fluorescent seeds. Single-color seeds, resulting from recombination within the interval, are selected. DNA is extracted from the resulting plants, and the target interval is amplified using long-range PCR (LR-PCR). The LR-PCR products are then sequenced to precisely identify crossover breakpoints. Data from multiple recombinant individuals are used to generate a high-resolution crossover map of the analyzed interval. **b.** Crossover landscape within *ChP* for theCol-*ChP* JMJ14 × L*er* (red) compared to the wild type (Col-*ChP* × L*er*, blue). Crossover frequencies are normalized to the total *ChP* crossover frequency. SNPs spaced at least 100bp apart (gray vertical lines) were used to determine the CO topology. The x-axis marks Col/L*er* SNPs, while red tick marks denote InDels. Genes are shown as light-green (forward strand) and violet (reverse strand) rectangles at the top; transposons are shown as orange rectangles. Positions of the three crossover hotspots (*Aro*, *Coco* and *Nala*) are highlited with colored rectangles below the plot. gRNAs used for JMJ14 targeting are indicated with burgundy arrowheads. Wild type data from Szymanska-Lejman et al. (2023). **c.** Genotypes of 164 Col-*ChP* JMJ14 × L*er* recombinants. Each horizontal line represents a single recombinant with color-coded genotypes along the x-axis (coordinates within *ChP*). **d.** Topological changes in crossover frequency across *ChP* caused by JMJ14 targeting to *Coco*, plotted as differences from the wild type. *Aro*, *Coco*, and *Nala* hotspots are highlighted with colored rectangles. The dashed blue horizontal line indicates the wild-type crossover level. gRNAs for JMJ14 are marked with burgundy arrowheads. Four sectors within *Coco* showing differential crossover remodeling are labeled *a–d*. **e.** Hotspot usage (in cM) for Col-ChP JMJ14 × Ler compared to Col-*ChP* × L*er*. **f.** Same as in **c**, but for Col-*ChP* × L*er*Δ#24 cross (data from Szymanska-Lejman et al., 2023). The Δ#24 deletion within *Coco* is marked by the transparent red rectangle. gRNA positions used for JMJ14 targeting are shown as burgundy lines for comparison. **g.** Hotspot usage (in cM) for Col-*ChP* × L*er*Δ#24 compared to Col-*ChP* × L*er*.

Comparison of crossover distribution in lines with dCas9-JMJ14 targeted to the *Coco* hotspot revealed significant differences from the wild type (Fig. 4b,c). We observed changes in the recombination topography of the *Coco* hotspot, reflected in a markedly altered crossover profile. Specifically, within the region targeted by JMJ14, there was a clear reduction in crossover frequency. However, immediately downstream of this region – still within the *Coco* interval – a short sector displayed an increase in crossover frequency.

To better characterize these topological differences, we performed a differential analysis in which the crossover distribution was normalized to that of the wild type (Fig. 4d). This analysis revealed that the reduction in crossover frequency occurred in two sectors (*b* and *d* in Fig. 4d), separated by a narrow sector (*c*) showing increased crossover frequency. Additionally, a rise in crossover frequency was observed in the distal right-hand sector (*a*). Interestingly, a reduction in crossover frequency was also apparent across the other two hotspots, *Aro* and *Nala* (Fig. 4d), which is further reflected in the hotspot usage analysis (Fig. 4e).

To further investigate the nature of the changes observed in JMJ14-targeted lines, we compared them to changes at the *ChP* interval in lines carrying a deletion of the *Coco* region. For this, we used our previously published data from the Col-*ChP* × L*er*Δ#24 cross (Szymanska-Lejman *et al*, 2023), normalizing the crossover distribution to that of the Col-*ChP* × L*er* wild type (Fig. 4f). The L*er*Δ#24 deletion lies within the same region as the targeted area, although it is slightly smaller than the region covered by the four gRNAs used for JMJ14 targeting (Fig. 4f). Despite this, none of the sectors within *Coco* exhibited an increase in crossover frequency relative to the wild type in the Col-*ChP* × L*er*Δ#24 cross. Consequently, the overall reduction in crossover frequency within *Coco* for this cross was greater than that observed in the Col-*ChP* JMJ14 × L*er* cross (80.3% versus 47.2%; Fig. 4g). In contrast, the other two hotspots showed virtually no reduction in crossover frequency, which differs markedly from the JMJ14-targeted lines (Fig. 4e and g, and Appendix Fig. S6).

In summary, targeting JMJ14 to a recombination hotspot leads to a local reduction in crossover frequency in sectors directly targeted by the gRNAs. However, the hotspot is not entirely silenced, as a fraction of DSBs is still repaired via crossover in the non-targeted sectors. At the same time, the inhibitory effect of JMJ14 is also visible in neighboring, non-targeted hotspots, suggesting a local spreading of the effector.

### Targeting JMJ14 to other chromosomal regions reduces their local crossover frequency

Conclusions drawn from a single genomic interval are subject to considerable uncertainty, particularly that the *ChP* interval is located relatively close to the centromere, and the *Coco* hotspot is one of the strongest recombination hotspots in *A. thaliana*, making it unique. To assess the broader applicability of the CRISPR–dCas9–JMJ14 system for reducing meiotic recombination via targeting H3K4me3 demethylation, we extended our approach to two additional genomic intervals. Specifically, we generated two additional reporter lines: *End3a*, a 413.7 kb interval located in a subtelomeric region containing the previously described hotspot *3a* (Yelina *et al*, 2012; Choi *et al*, 2013), and *Crimson Wave* (*CW*), a 72.3 kb interval located in the middle of a chromosome arm (Fig. 5a and Appendix Figs. S7 and S8). Details on the construction of *End3a* and *CW* lines are provided in the Appendix Fig. S9 and in the Methods section.

**Figure 5.**
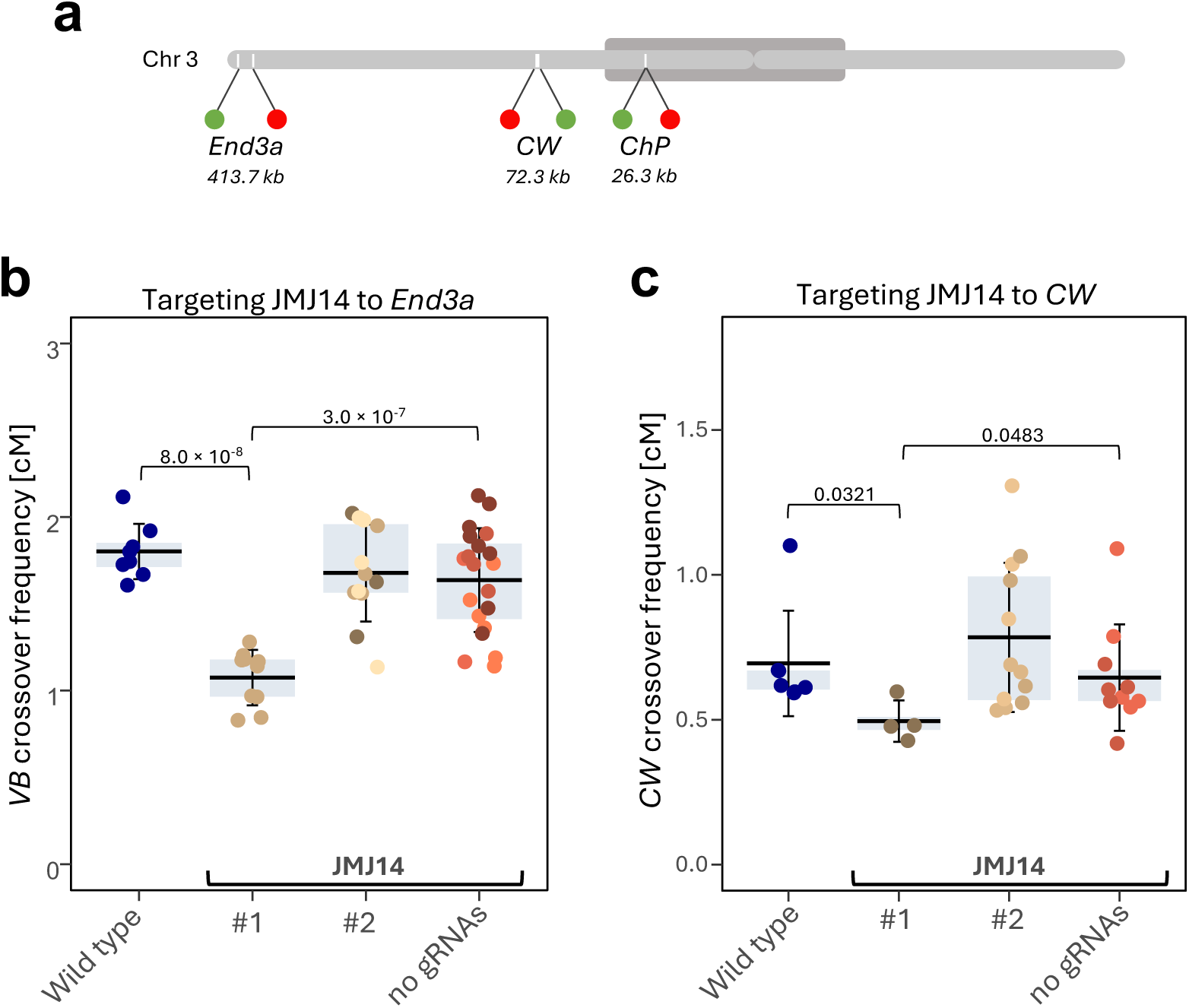
Targeting JMJ14 reduces crossover hotspot activity independently of chromosomal location. **a.** Schematic of *A. thaliana* chromosome 3, highlighting the *End3a*, *CW*, and *ChP* intervals. Red and green markers indicate the positions of dsRed and eGFP reporters, while the dark gray oval represents the pericentromeric region. **b.** Crossover frequency in the subtelomeric *End3a* interval for crosses with transgenic lines expressing dCas9-JMJ14 targeted to the *3a* hotspot (#1-3). Controls include wild-type Col-*End3a* × L*er* crosses and transgenic lines expressing dCas9-JMJ14 without gRNAs. In the boxplot, the centre line represents the median, the box spans the interquartile range (25th–75th percentile), and whiskers indicate the standard deviation. Each data point represents the crossover frequency of a single cross. Statistical significance was assessed using Welch’s test. **c.** As in (**b**), but for the interstitial *CW* interval.

As in previous experiments, after transforming plants with the construct, individuals carrying the transgene were crossed with wild-type L*er* plants to measure crossover frequency based on reporter segregation in F₂ seeds. In the *End3a* line, one of the three analyzed T₁ lines exhibited a significant, nearly twofold reduction in crossover frequency compared to wild type (Fig. 5b; *P* = 7.95e-8, Welch’s test). Similarly, in the *CW* line, one of three analyzed lines showed a significant reduction in crossover frequency (Fig. 5c; *P* = 0.0321, Welch’s test).

These observations indicate that targeting JMJ14 using a CRISPR-dCas9-based system can locally reduce meiotic recombination at any chosen hotspot, regardless of its chromosomal location. However, not all transgenic lines carrying the dCas9 construct were equally effective. Such variability is a common characteristic of genetic constructs integrated into the host genome via *Agrobacterium*-mediated transformation and is generally attributed to insertion site effects and construct expression levels (Jupe *et al*, 2019). Additionally, the observed reduction in crossover frequency in *no gRNA* lines suggests that, beyond the localized effect achieved through precise CRISPR-gRNA targeting, a portion of the dCas9-JMJ14 fusion protein may act non-specifically at random genomic locations.

### Local transcriptional activity and associated chromatin changes induced by VP64 can stimulate crossover recombination

Modulating H3K4me3 levels by targeting JMJ14 resulted in both a decrease in crossover frequency and a reduction in the expression of the gene encoding *CocoRNA*. Therefore, we asked whether local induction of gene expression could increase crossover frequency in this interval. To address this, we decided to use the synthetic activation domain VP64, a derivative of the herpes simplex virus (HSV-1) activation domain (Sadowski *et al*, 1988; Beerli *et al*, 1998), targeted to the *CocoRNA* promoter via translational fusion with dCas9. VP64 is a universal transcriptional activator that functions in both animals and plants (Lowder *et al*, 2015; Cano-Rodriguez *et al*, 2016; Lowder *et al*, 2018). It interacts with transcriptional mediators and RNA polymerase II, enhancing transcription initiation when delivered near the transcription start site (TSS) (Kim *et al*, 1994; Allen & Taatjes, 2015).

Since *CocoRNA* encodes a lncRNA that had not been extensively characterized, we first sought to identify its TSS using 5’ RACE in Col and L*er*. The TSS was identical in both accessions (Appendix Fig. S10), and based on this information, we designed three gRNAs located +30 bp, +234 bp, and +477 bp upstream of the TSS. We then generated transformants expressing dCas9-VP64 targeted by these three gRNAs in the Col-*ChP* background (Appendix Fig. S11a). Despite high dCas9-VP64 expression in all T_1_ lines (Appendix Fig. S11b), only one line exhibited a threefold increase in *CocoRNA* expression levels (Fig. 6a). This single line was used for crosses with wild-type Col to measure local recombination rates. In the F_1_ generation, we observed an increase in *ChP* crossover frequency from 0.33 cM in wild-type to 0.52 cM (Fig. 6b; *P* = 6.7 × 10⁻⁴, Mann-Whitney U test).

**Figure 6.**
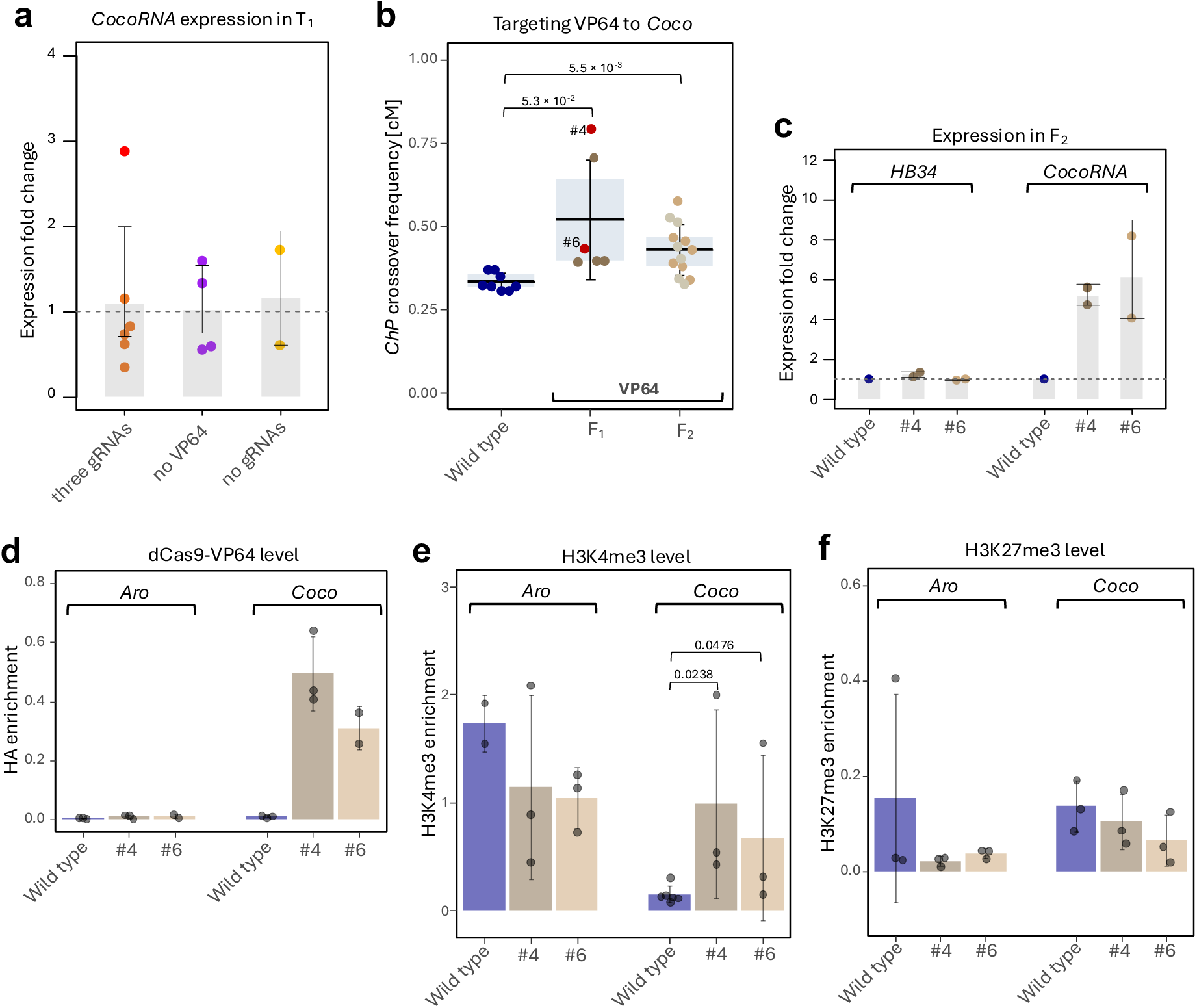
Targeting VP64 enhances crossover hotspot activity. a. Relative expression of the the *CocoRNA* gene within the *Coco* hotspot in T_1_ plants carrying the dCas9-VP64 construct targeted to the 5’ UTR of *CocoRNA*, measured by RT-qPCR. Bars represent the mean values of biological replicates (dots), with error bars indicating standard deviation. Each dot represents an invdividual T_1_ plant. A single plant exhibiting elevated expression is highlighted in red. b. Crossover frequency in the *ChP* interval in crosses with the T_1_ plant exhibiting elevated CocoRNA expression (highlighted in red in (**a**)) in the F_1_ cross and F_2_ generation. Each data point represents a measurement for an individual plant. OQspring from F_1_ individuals #4 and #6 are shown in a single graph, distinguished by brown and gray colors, respectively (F_2_). Statistical significance was assessed using Welch’s test. c. Expression levels of *HB34* (*Aro* hotspot) and *CocoRNA* (*Coco* hotspot) in F_2_ plants, measured by RT-qPCR. Bars represent the mean values of two biological replicates (dots), with error bars indicating standard deviation. Dashed line indicates the wild type level. d. Enrichment specificity of dCas9-VP64 at the *Aro* (non-targeted) and *Coco* (targeted) hotspots in F_2_ offspring from plants #4 and #6, measured by ChIP-qPCR. Bars represent the mean values of biological replicates (dots), with error bars indicating standard deviation. e. Changes in H3K4me3 enrichment at the *Aro* (non-targeted) and *Coco* (targeted) hotspots in F_2_ offspring from plants #4 and #6, measured by ChIP-qPCR. Bars represent the mean values of biological replicates (dots), with error bars indicating standard deviation. Statistical significance was assessed using Mann-Whitney U test. f. As for e, but for H3K27me3 enrichment.

Since these transformants were crossed with Col-background plants, self-fertilization was possible, allowing us to directly analyze the F_2_ generation. We selected progeny from two F_1_ plants differing in crossover frequency. RT-qPCR confirmed that both families of F_2_ plants exhibited similarly elevated *CocoRNA* expression compared to wild type (Fig. 6c). Importantly, expression of *HB34*, a gene located in the adjacent *Aro* hotspot, remained unchanged, indicating high specificity of transcriptional activation by dCas9-VP64 (Fig. 6c). This was consistent with strong and *Coco*-specific enrichment of dCas9-VP64 (Fig. 6d).

Furthermore, ChIP-qPCR analysis showed that F_2_ plants exhibited nearly up to threefold increase in H3K4me3 levels in the *Coco* region compared to wild type (Fig. 6e, *P* < 0.048, Mann-Whitney U test). As expected, no enrichment was observed for H3K27me3, a repressive chromatin mark (Fig. 6f). Crossover frequency in the *ChP* interval in the F_2_ generation was significantly higher than in wild-type plants. However, unlike in the F_1_ generation, no difference was observed between the two F_2_ lines (Fig. 6b).

In summary, targeting VP64 to the *Coco* recombination hotspot leads to chromatin modifications associated with open chromatin structure, transcriptional activation of the gene located within the hotspot, and a local increase in crossover frequency.

## DISCUSSION

EQorts to directly modulate recombination hotspot activity in plants are substantially limited by the inherently low crossover frequency and the presence of a vast number of relatively weak hotspots. A single chromosome may contain thousands of promoters and gene terminators, each of which can serve as a potential recombination hotspot (Choi *et al*, 2013). However, only one to three crossover events typically occur per chromosome during a single meiosis. Thus, detecting changes in the activity of individual hotspots requires highly specialized approaches. In this study, we utilized a recently developed reporter system that enables the measurement of recombination within short chromosomal intervals containing only a few recombination hotspots. By analysing reporter segregation across thousands of seeds, we could detect subtle changes in crossover frequency within a defined interval, thereby assessing the local impact of targeted chromatin modifications on individual hotspots.

Our main objective was to determine which chromatin modifiers are capable of inducing local, targeted changes in recombination activity. To this end, we employed effector domains translationally fused to dCas9 and guided to specific genomic loci using three or four gRNAs. The effectors included catalytic domains of histone methyltransferases KYP and CLF, which deposit repressive chromatin marks (H3K9me2 and H3K27me3, respectively), as well as methyltransferases ATX2, SDG4, SDG8, and PRDM9, which deposit activating marks such as H3K4me2/3 and H3K36me3 (Jackson *et al*, 2002; Schubert *et al*, 2006; Saleh *et al*, 2008; Cartagena *et al*, 2008; Kim *et al*, 2005; Zhao *et al*, 2005; Baudat *et al*, 2010; Myers *et al*, 2010; Parvanov *et al*, 2010). We also tested the demethylase JMJ14, which removes H3K4me2/3, and the general transcriptional activator VP64 (Lu *et al*, 2010; Yang *et al*, 2010; Lowder *et al*, 2015; Cano-Rodriguez *et al*, 2016). In total, we evaluated the effects of eight different chromatin-modifying effectors.

Notably, none of the tested histone methyltransferases caused a significant change in crossover frequency. For SDG4 and SDG8, and to a lesser extent ATX2, a major complication was the strong silencing of the endogenous genes encoding these enzymes, which was observed already in the T_1_ generation. Recent work has shown that this silencing effect can be alleviated by using an *rdr6* mutant background (Wang *et al*, 2023). Targeting the related methyltransferase SDG2 in such a background resulted in elevated crossover frequency within pericentromeric regions (Binenbaum *et al*, 2025). Interestingly, in that study, the increase in H3K4me3 enrichment was not localized precisely to the targeted centromeric regions but was instead detected in their flanking domains, and to a lesser extent, throughout the genome. In contrast, targeting PRDM9 – a methyltransferase absent from the plant lineage – had no detectable effect on crossover frequency (Fig. 2c). Collectively, these findings suggest that while histone methyltransferases have the potential to modulate meiotic recombination, their practical use in targeted applications is limited by the risk of oQ-target effects, particularly the silencing of endogenous chromatin regulators.

In contrast, targeting the demethylase JMJ14 to recombination hotspots proved effective. In crosses involving T_1_ plants expressing dCas9-JMJ14 constructs, we observed up to a twofold reduction in crossover frequency within the *ChP* interval, despite targeting being confined to the central region of the *Coco* hotspot (Fig. 3a). These changes were associated with a reduction in H3K4me3 enrichment and decreased expression of *CocoRNA*, a long non-coding RNA gene located at the centre of the *Coco* hotspot (Fig. 3c,d). Interestingly, the neighboring *Aro* hotspot gene *HB34* exhibited increased transcription, suggesting coregulation of these loci (Fig. 3d). It is worth noting that JMJ14 was recently identified as one of the few effector proteins capable of inducing targeted gene repression, supporting its utility in chromatin remodelling applications (Wang *et al*, 2023).

JMJ14 targeting was carried out using four gRNAs directed to the hyperactive central region of *Coco*, the strongest hotspot within the *ChP* interval. By sequencing 164 recombinants through a seed-typing approach, we were able to precisely assess the effects of JMJ14. Beyond the reduction in crossover frequency in the targeted region, we observed increases in recombination in neighbouring regions of the same hotspot. This pattern contrasts with results from lines carrying a deletion of the hyperactive region of *Coco*, where a reduction in crossover frequency was not accompanied by compensatory increases elsewhere in the hotspot. These results indicate that JMJ14-mediated demethylation of H3K4me2/3 does not fully suppress crossover formation but instead leads to a partial redistribution of crossovers within the hotspot.

We also observed a reduction in crossover frequency at the adjacent *Aro* and *Nala* hotspots in lines with JMJ14 targeted to *Coco*, likely contributing to the overall reduction in recombination within the *ChP* interval. This effect was absent in the deletion line, in which the flanking hotspots remained unaffected. These results imply that JMJ14-mediated demethylation spreads beyond the directly targeted regions – a phenomenon commonly associated with dCas9-based systems (Thakore *et al*, 2016).

Since our attempts to increase H3K4me3 levels by targeting histone methyltransferases did not yield the expected recombination effects, and JMJ14 targeting also reduced the expression of the targeted gene, we explored the use of a general transcriptional activator, VP64, to enhance hotspot activity. This approach proved successful: in crosses involving T_1_ lines expressing dCas9-VP64 and showing elevated expression of the targeted gene, we observed a significant increase in crossover frequency within the *ChP* interval (Fig. 6). VP64 is known to recruit histone acetyltransferases and transcriptional co-activator complexes, simultaneously affecting chromatin structure and transcription (Uesugi *et al*, 1997; Perez-Pinera *et al*, 2013; Chavez *et al*, 2015). It remains unclear to what extent the increased hotspot activity is driven by enhanced *CocoRNA* transcription versus VP64-associated chromatin modifications.

In summary, our experiments demonstrated that modification of H3K4me2/3 affects meiotic recombination and can be used for local, targeted changes in crossover frequency via dCas9-mediated targeting. We also showed that epigenetic factors promoting transcriptional activation can simultaneously lead to an increase in crossover frequency. At the same time, it appears that the chromatin changes we induced do not fully explain the differences in the activity of particular recombination hotspots.

## Methods

### Growth conditions and plant material

Plants were cultivated under controlled conditions, including a temperature of 22 °C, 60-70% relative humidity, 150-μmol light intensity and a 16-hour light/8-hour dark photoperiod. Seeds were stratified at 4°C for 48 hours before transferring to growth chambers.

Fluorescence-tagged lines (FTLs) used in this study included *End3a*, *Crimson Wave* (*CW*), and *Chili Pepper* (*ChP*). While *ChP* had been previously characterized (Szymanska-Lejman *et al*, 2023), the *End3a* and *CW* lines were generated by crossing two single-colour lines carrying coding sequences of the seed-expressed fluorescent transgenes (eGFP or dsRed). The single-color FTLs were kindly provided by Dr. Scott Poethig. Specifically, *CG821* and *CR921* were crossed to generate *End3a*, and *CR729* and *CG739* were crossed to obtain the *CW* interval. The exact locations and sizes of the intervals are shown in Appendix Fig. S9. F_1_ progeny were self-fertilized, and F_2_ seeds were screened using a Zeiss Lumar V12 epifluorescent stereomicroscope. Crossover (CO) events between reporter genes were identified based on fluorescence patterns, specifically seeds displaying a homozygous state for one reporter and a hemizygous state for the other (GR/–R or GR/G–) were preselected. These F₂ plants were subsequently selfed, and intervals with homozygous fluorescent reporters were selected for further study.

### Recombination frequency measurements

Recombination frequencies in the obtained FTLs was assessed using a previously described seed-based fluorescence system (Szymanska-Lejman *et al*, 2023). Seeds were collected from hemizygous *End3a*, *CW*, or *ChP* (RG/--) plants and cleaned via sieve filtration. Images of F_2_ seeds were acquired using a Zeiss epifluorescent stereomicroscope, capturing brightfield and fluorescence images through red and green filters. Seed counts were performed using a custom in-house Python script. Due to the low recombination frequencies, particularly in the CW and ChP lines, single-color recombinant seeds were manually identified. Recombination frequency (RF), expressed in centiMorgans (cM), was calculated using the formula: RF = 100 × (1 – (1 – 2(NG + NR)/NT)/2) where *NG* is the number of green-only seeds, *NR* is the number of red-only seeds, and *NT* is the total number of seeds collected per plant. Raw seed scoring data for all measurements are provided in the Appendix Data.

### Preparation of sgRNA expression cassettes

The single guide RNA (sgRNA) expression cassettes were prepared as previously described (Bieluszewski *et al*, 2022b). gRNAs targeting hotspots within *ChP*, *End3a* or *CW* intervals were designed using an available tool, CRISPOR (http://crispor.tefor.net). All gRNA spacer sequences and their corresponding genomic coordinates are listed in the Appendix Table S1.

### Constructs preparation and plant transformation

Expression cassettes were amplified with CloneAmp high fidelity polymerase (Takara). Histone methyltransferase and demethylase domains were amplified from cDNA, while VP64 and dCas9-HA coding sequences were obtained from a previously described SunTag plasmid (Papikian *et al*, 2019). All primers used for cloning are provided in the Appendix Table S2. sgRNA expression cassettes were cloned following the method described by Bieluszewski *et al*, (2022b). Genetic constructs were assembled using the Gibson assembly method (ClonExpress MultiS One Step Cloning Kit II, Vazyme), with effector domains inserted in-frame between the dCas9 and HA-tag sequences.

Prepared binary constructs were introduced into *Agrobacterium tumefaciens* cells and used for plant transformation via floral dipping as described (Bieluszewski *et al*, 2022b). T_1_ transformants were selected either by applying BASTA herbicide or by genotyping for construct presence (primers listed in the Appendix Table S2).

### ChIP-qPCR

Chromatin was isolated from 2 g of *Arabidopsis* leaves with use of Honda buffer (440 mM sucrose, 25 mM Tris-HCl pH 8.0, 10 mM MgCl_2_, 0.5 % Triton X-100, 10 mM β-ME, Roche Protein Inhibitor Cocktail, 10 mM PMSF, 40 mM spermine). Chromatin was pre-cleared for 1 h with Dynabeads Protein A (Thermo Fisher Scientific) and then immunoprecipitation was carried out through overnight incubation in 4°C with 2-3 ng of selected antibodies (α-HA C29F4 Cell Signalling Technology, α-H3K4me3 ab8580, α-H3K4me3 04-745 Merck, α-H3K27me3 ab6002 Abcam, α-H3 ab1791 Abcam). To capture immunocomplexes, Dynabeads Protein A were added and incubated for 1 h in 4°C. Then, beads were washed twice with low-salt (150 mM NaCl), high-salt (500 mM NaCl), LiCl (10 mM Tris-HCl, pH 8.0, 1 mM EDTA, 0.25 M LiCl, 1% Nonidet P-40, 0.5% sodium deoxycholate, 1 mM PMSF) and TE buffers. Finally, complexes were eluted by incubation with 10% Chelex in 99°C. Following treatment with proteinase K, the resulting ChIP-DNA was diluted and used as a template in qPCR reactions. Percent of input for each tested histone modifications was normalized to Input DNA.

### RT-qPCR

30 mg of unopened flower buds were collected and used to isolate RNA with use of RNeasy Plant Mini Kit (Qiagen). cDNA synthesis was performed using 1 μg of total RNA and oligo-dT primers (HiScript III, Vazyme). cDNA was diluted fivefold and used as a template in RT-qPCR reactions (SYBR Green qPCR Master Mix, Thermo Fisher Scientific). Two reference genes (*AT3G18780* and *AT1G14400*) were used as controls.

## 5’RACE

The RNA from ∼30 mg unopened flower was isolated with RNeasy Plant Mini Kit (Qiagen). 1 µg of RNA was used to generate cDNA with oligo-dT and random hexamers (HiScript III, Vazyme). 5’RACE kit (Roche) was used to capture the upstream regions of lncRNA by subsequent nested PCR reactions. Obtained amplicons were purified and cloned into pJET1.2 (Thermo Fisher Scientific) plasmid which was later transformed into competent DH5α *Escherichia coli* strain. Colony PCR with insert-specific and flanking primers was used to screen for clones with specific insert. Eight independent plasmid clones were sequenced for both Col and L*er*.

### Seed-typing for the *ChP* Interval

The seed-typing method for fine-scale mapping of crossover (CO) breakpoints was previously described by Szymanska-Lejman *et al.,* (2023). Briefly, recombinant seeds were manually preselected from F_2_ *Col-ChP × Ler* plants carrying *dCas9-JMJ14* targeted to the *Coco* hotspot. DNA was extracted using a CTAB buffer and purified with AMPure XP magnetic beads (Beckman-Coulter) to ensure high-quality templates essential for long-range PCR (LR-PCR). LR-PCR was performed using PrimeSTAR GXL Polymerase (TaKaRa Bio) under the following conditions: 0.2–10 ng of template DNA, 1.2 µL of 2.5 mmol/L primers, 3 µL buffer, 1.2 µL dNTPs, 0.3 µL polymerase, and distilled water to a final volume of 15 µL. The cycling conditions were 30 cycles at 98°C for 10 s and 68°C for 10 min. The *ChP* interval was amplified in three separate reactions, generating 9–11 kb amplicons, which were visualized by gel electrophoresis.

Amplicons from the same recombinant sample were pooled and purified with magnetic beads for library preparation, performed as previously described (Szymanska-Lejman *et al*, 2023). Briefly, 1 µL of pooled PCR products was tagmented with Tagmentation Buffer (40 mM Tris-HCl pH 7.5, 40 mM MgCl₂), 0.5 µL DMF (Sigma), 2.35 µL nuclease-free water (Thermo Fisher), and 0.05 µL in-house loaded Tn5 transposase. The reaction was incubated at 55°C for 2 min and stopped with 1 µL 0.1% SDS, followed by a 10 min incubation at 65°C. Tagmented DNA was then amplified using the KAPA2G Robust PCR kit (Sigma) with unique P5/P7 indexing primers (Rowan *et al*, 2019). Indexed libraries were pooled and sequenced on an Illumina NovaSeq platform. CO breakpoints were identified based on single-nucleotide polymorphisms (SNPs) differentiating the two parental accessions.

Reads from libraries prepared from 164 recombinants preselected from F_2_ Col-*ChP* × *Ler* plants carrying *dCas9-JMJ14* targeted to the *Coco* hotspot were aligned to genomic sequence of *ChP* interval with bwa-mem (Li & Durbin, 2009). Resulting bam files were sorted and indexed with samtools (Li, 2011). Polymorphic sites were called against previously generated high-fidelity SNP list obtained from seed-typing of 243 individual F_2_ Col-*ChP* × L*er* plants (Szymanska-Lejman *et al*, 2023). SNPs with associated reads were used to call haplotype blocks and denominate crossover sites across the *ChP* interval. Data were visualized using a publicly available custom R script (https://github.com/LabGenBiol/ESILs). Raw sequencing data of 164 recombinants preselected from Col-*ChP × Ler* plants carrying *dCas9-JMJ14* targeted to the *Coco* hotspot were deposited in GEO database under BioProject number PRJNA1254937.

## Acknowledgements

We thank Dr. Scott Poethig for kindly providing the single-color FTLs. This work was supported by the Foundation for Polish Science grant POIR.04.04.00-00-5C0F/17-00 to P.A.Z., and the National Science Centre, Poland (NCN) grants 2021/41/N/NZ2/00340 to M.S.-L. and 2020/39/I/NZ2/02464 to P.A.Z.

## Data availability

The datasets produced in this study are available in the following database: GBS data: Gene Expression Omnibus PRJNA1254937 (reviewer’s link: https://dataview.ncbi.nlm.nih.gov/object/PRJNA1254937?reviewer=239oquit12tjc613p3kbn30dfr)

## Author contributions

W.D., M.S.-L. and P.A.Z. designed research; W.D., M.S.-L., A.W. and K.H. performed research; W.D. performed the computational analyses; W.D., M.S.-L., A.W., K.H., and P.A.Z. analysed data; M.S.-L. and P.A.Z. wrote the paper.

## Appendix Figures

**Appendix Fig. S1.**
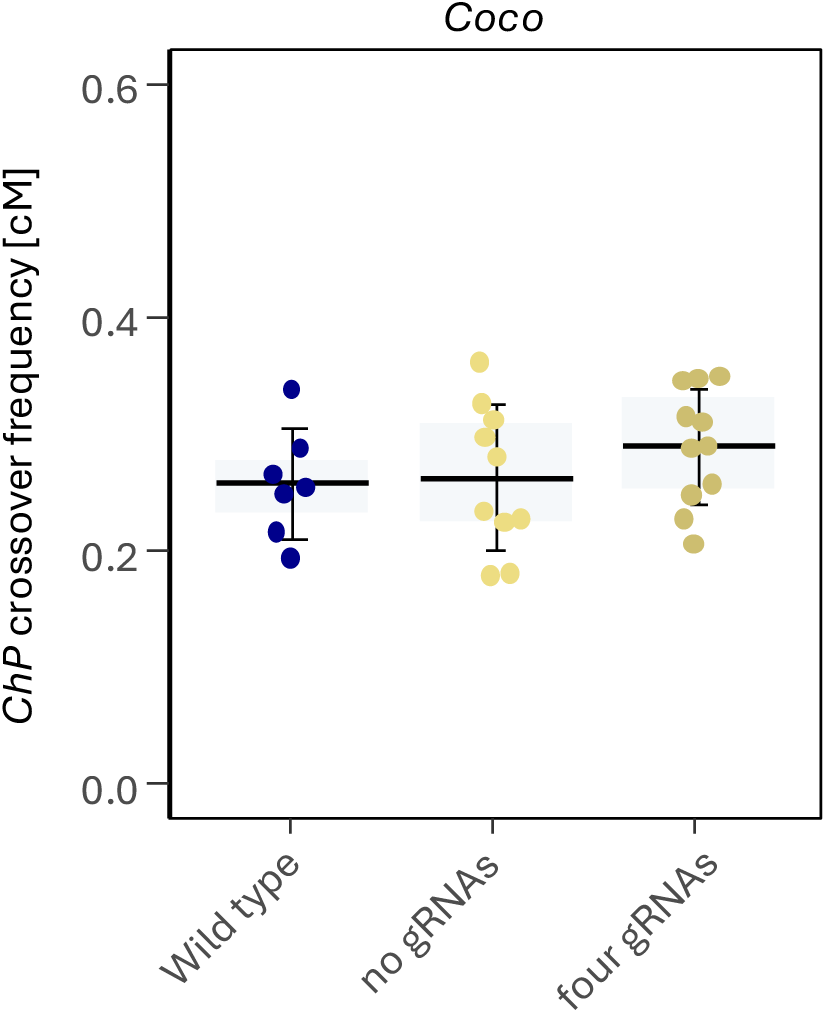
Crossover frequency in crosses with transformants expressing dCas9 without gRNAs or with four gRNAs targeting the *Coco* hotspot. The Col-*ChP* × Col cross served as the wild-type control. In the boxplot, the center line indicates the median, the upper and lower bounds represent the 75th and 25th percentiles, and error bars show the standard deviation. Each data point represents a crossover frequency measurement for a single cross. At least three independent transformants were used in each case.

**Appendix Fig. S2.**
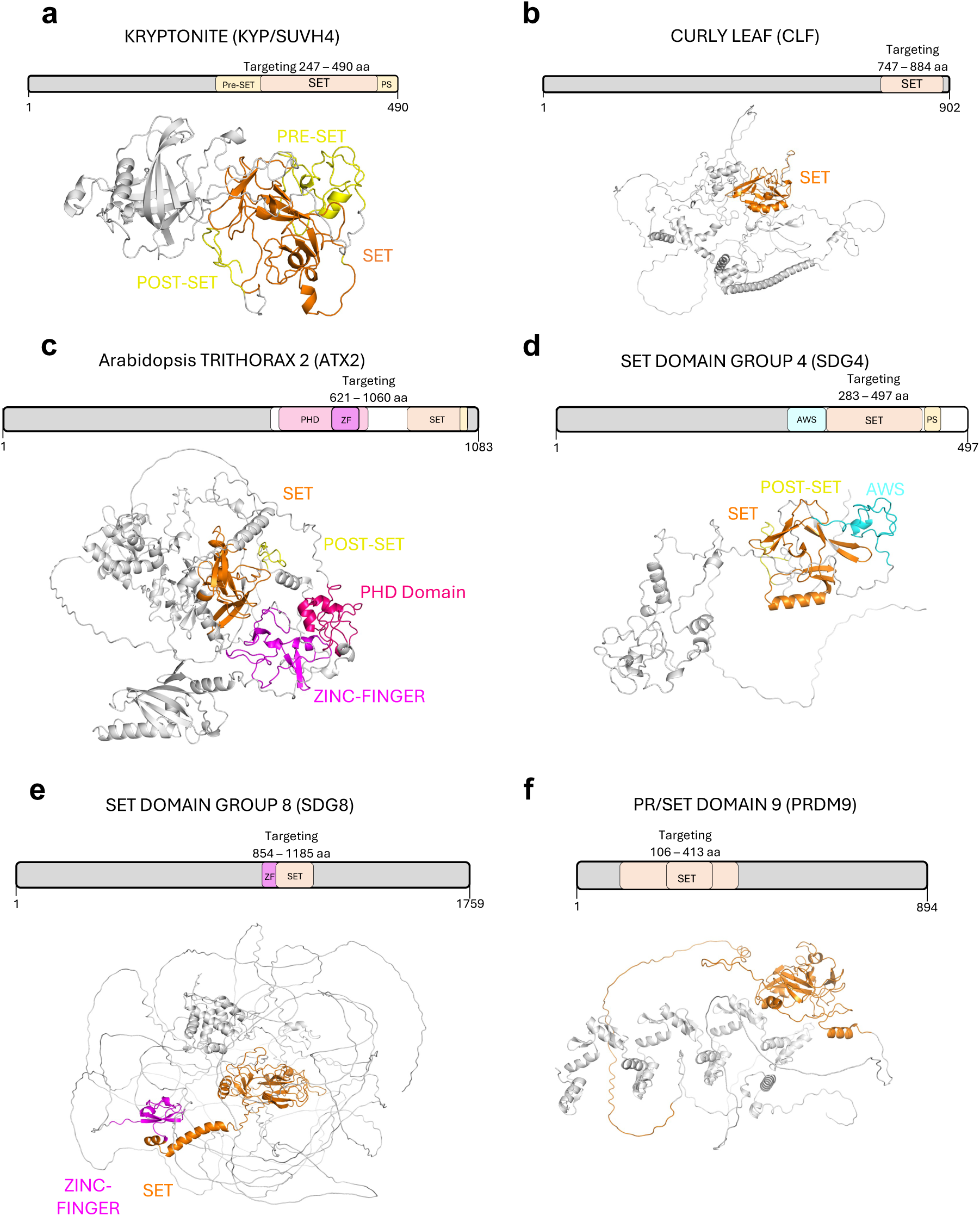
Domain architecture and predicted structure of the six histone methyltransferases used in the study. The upper panels depict the domain organization of each protein, with the regions used for translational fusions with CRISPR-dCas9 indicated. Key functional domains are highlighted: SET (Su(var)3-9, Enhancer-of-zeste, Trithorax domain), pre-SET, PS (post-SET domain), ZF (Zinc Finger), PHD (Plant Homeodomain), and AWS (Associated With SET domain). The lower panels present structural models of each protein predicted by AlphaFold2. The SET domain mediates methyltransferase activity by catalyzing the transfer of methyl groups from S-adenosylmethionine (SAM) to specific lysine residues on histone tail.

**Appendix Fig. S3.**
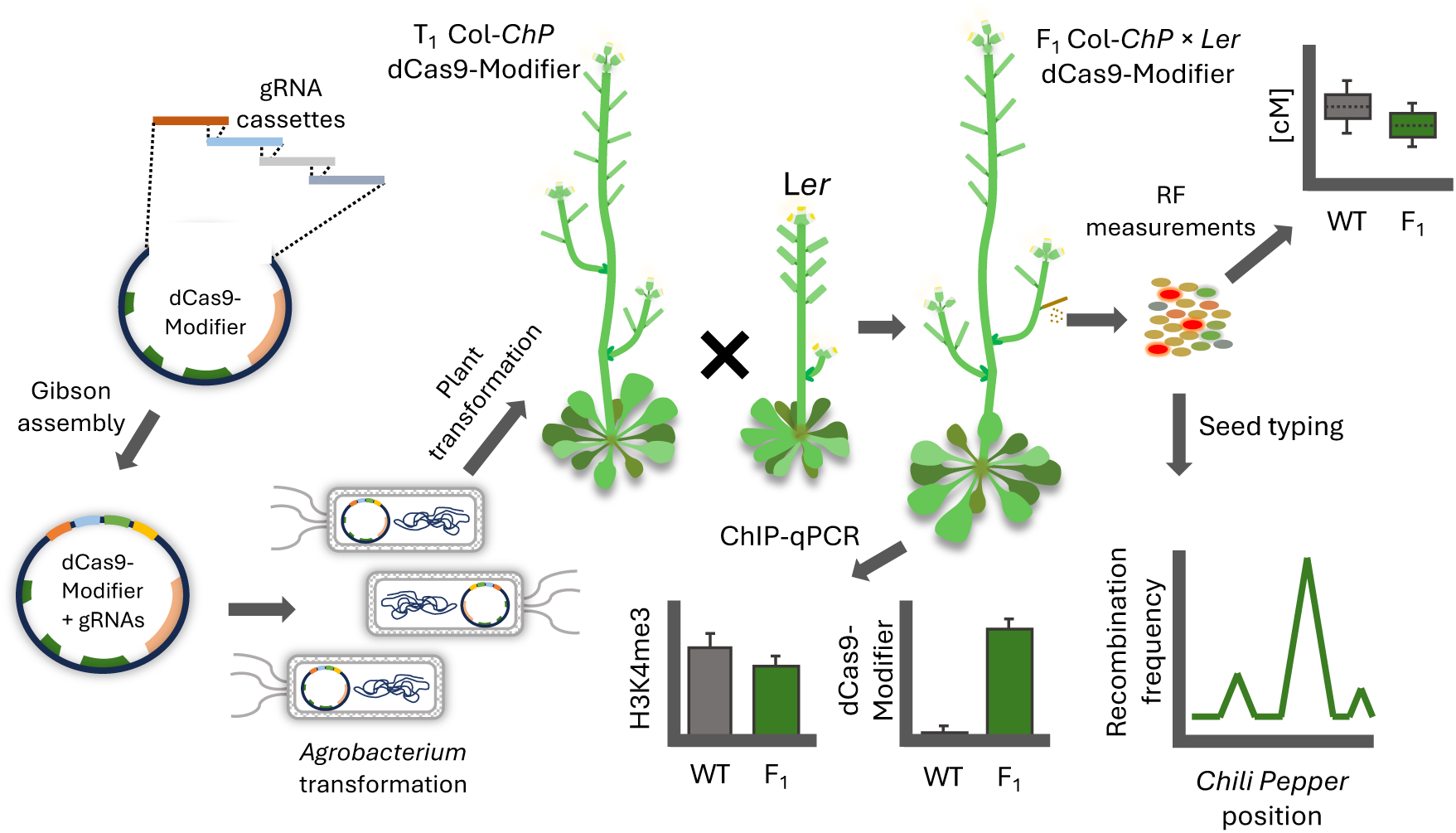
Experimental pipeline for screening H3 modifications affecting local crossover frequency. gRNAs targeting the *ChP* hotspot were cloned into a vector containing a translational fusion of dCas9 with a selected chromatin modifier and subsequently introduced into Col-*ChP* plants via *Agrobacterium*-mediated transformation. Alternatively, gRNAs targeting other hotspots were used to transform Col-*ChP*, Col-*End3a*, or Col-*CW* plants (not shown). The resulting transformants were crossed with L*er*, and the F_1_ hybrids were self-pollinated. The harvested seeds were used to measure recombination frequency (RF). If a change in RF was observed compared to wild-type control plants or plants transformed with constructs lacking gRNAs (not shown), H3K4me3 levels and dCas9 binding were analysed by ChIP-qPCR. In the case of the dCas9-JMJ14 line targeted to Col-*ChP*, a high-resolution crossover analysis was performed using the seed-typing technique.

**Appendix Fig. S4.**
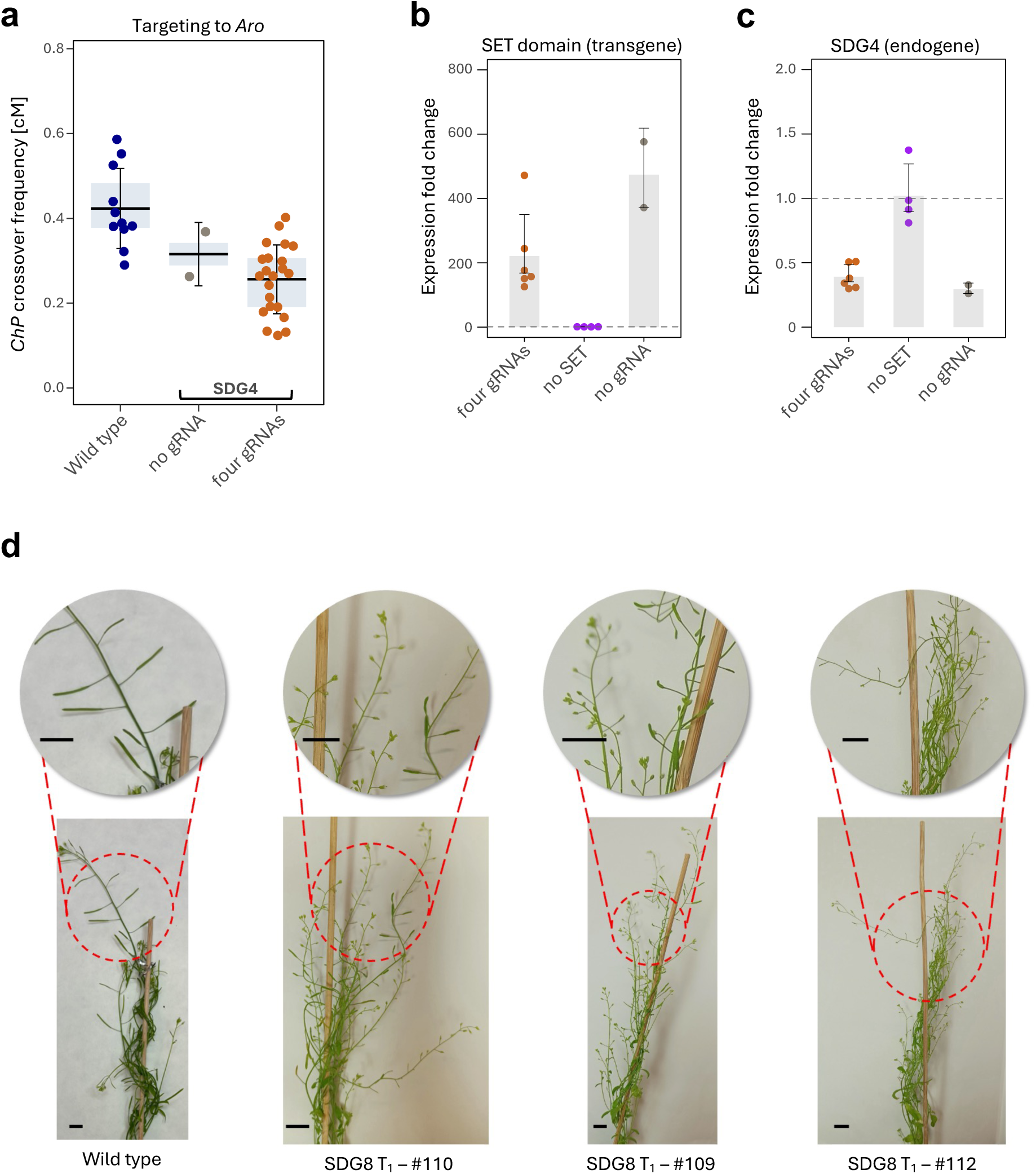
Silencing in lines targeting the SET domains of SDG4 and SDG8 to the *Aro* hotspot. a. Crossover frequency in the *ChP* interval in crosses with transformants carrying constructs expressing dCas9 fused to the SET domain of histone methyltransferases SDG4 targeted to the *Aro* hotspot using four gRNAs. Controls included plants without gRNAs and wild-type Col-*ChP* × L*er*. In the boxplot, the center line indicates the median, the upper and lower bounds represent the 75th and 25th percentiles, and error bars show the standard deviation. Each data point represents crossover frequency for a single cross. At least three independent transformants were analyzed per construct. b. Relative expression level of the SET domain in crosses with lines carrying the dCas9-SDG4 construct, with or without gRNAs, compared to wild-type plants, as determined by RT-qPCR. A control line expressing dCas9 with gRNAs (but without the SET domain) was included for comparison. Each dot represents a biological replicate (one cross). Bar plots show the average fold change relative to wild type, with error bars indicating standard deviation. Dashed line indicates wild type expression level. c. Relative expression level of the native *SDG4* gene in plants shown in (**b**). Each dot represents a biological replicate (one cross). Bar plots show the average fold change relative to wild type, with error bars indicating standard deviation. Dashed line indicates wild type expression level. d. T_1_ plants carrying the *dCas9-SET-SDG8* construct are sterile, most likely due to the silencing of the native *SDG8* gene. Inflorescences of three representative T_1_ plants are shown.

**Appendix Fig. S5.**
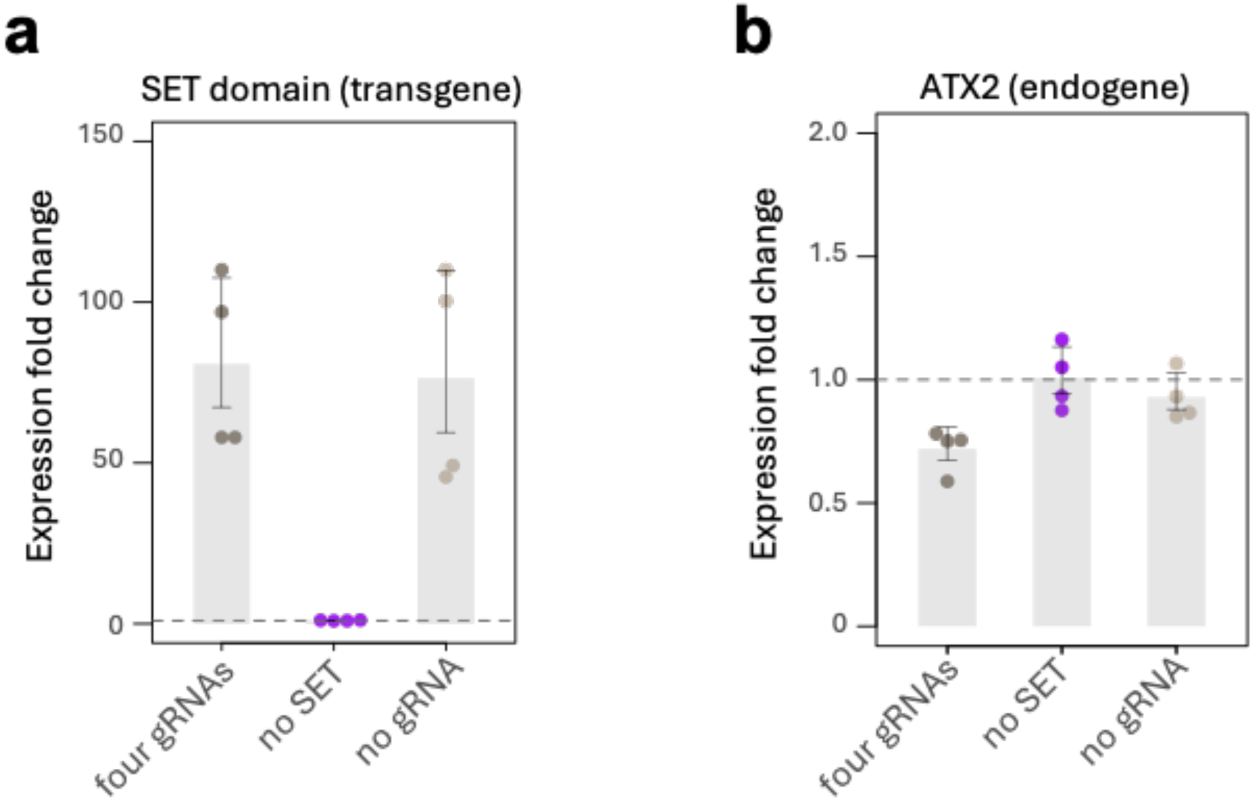
Partial silencing of *ATX2* expression in lines targeting the SET domains of ATX2 to the *Aro* hotspot. **a.** Relative expression level of the SET domain in crosses with lines carrying the dCas9-ATX2 construct, with or without gRNAs, compared to wild-type plants, as determined by RT-qPCR. A control line expressing dCas9 with gRNAs (but without the SET domain) was included for comparison. Each dot represents a biological replicate (one cross). Bar plots show the average fold change relative to wild type, with error bars indicating standard deviation. Dashed line indicates wild type expression level. **b.** Relative expression level of the native *ATX2* gene in plants shown in (**a**). Each dot represents a biological replicate (one cross). Bar plots show the average fold change relative to wild type, with error bars indicating standard deviation. Dashed line indicates wild type expression level.

**Appendix Fig. S6.**
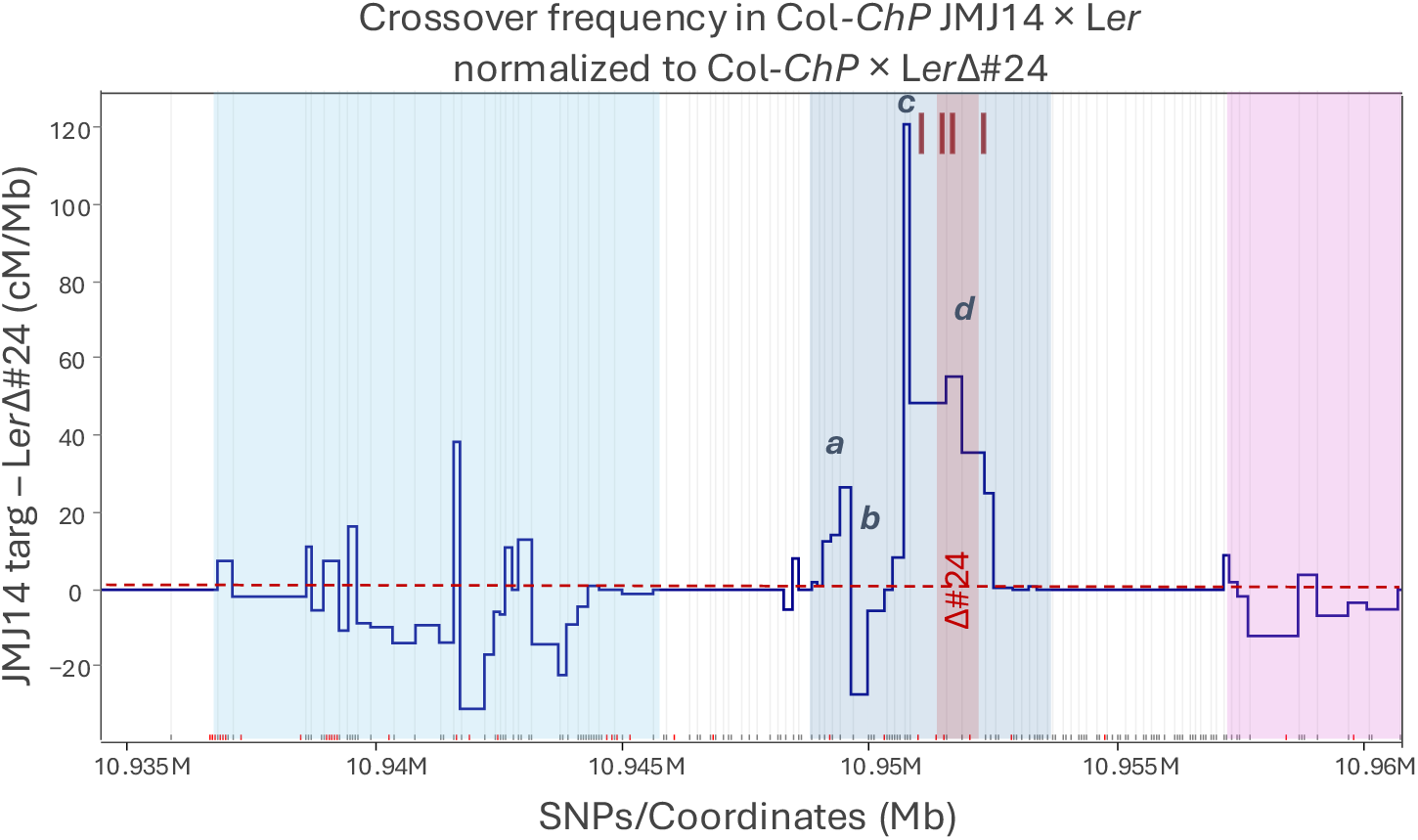
Topological changes in crossover frequency across *ChP* caused by JMJ14 targeting to *Coco*, plotted as differences from the Col-*ChP* × L*er*Δ#24 cross (data from Szymanska-Lejman et al., 2023). *Aro*, *Coco*, and *Nala* hotspots are highlighted with colored rectangles. The dashed red horizontal line indicates the Col-*ChP* × L*er*Δ#24 crossover level. gRNAs for JMJ14 are marked with burgundy lines. Four sectors within *Coco* showing differential crossover remodeling are labeled *a–d*.

**Appendix Fig. S7.**
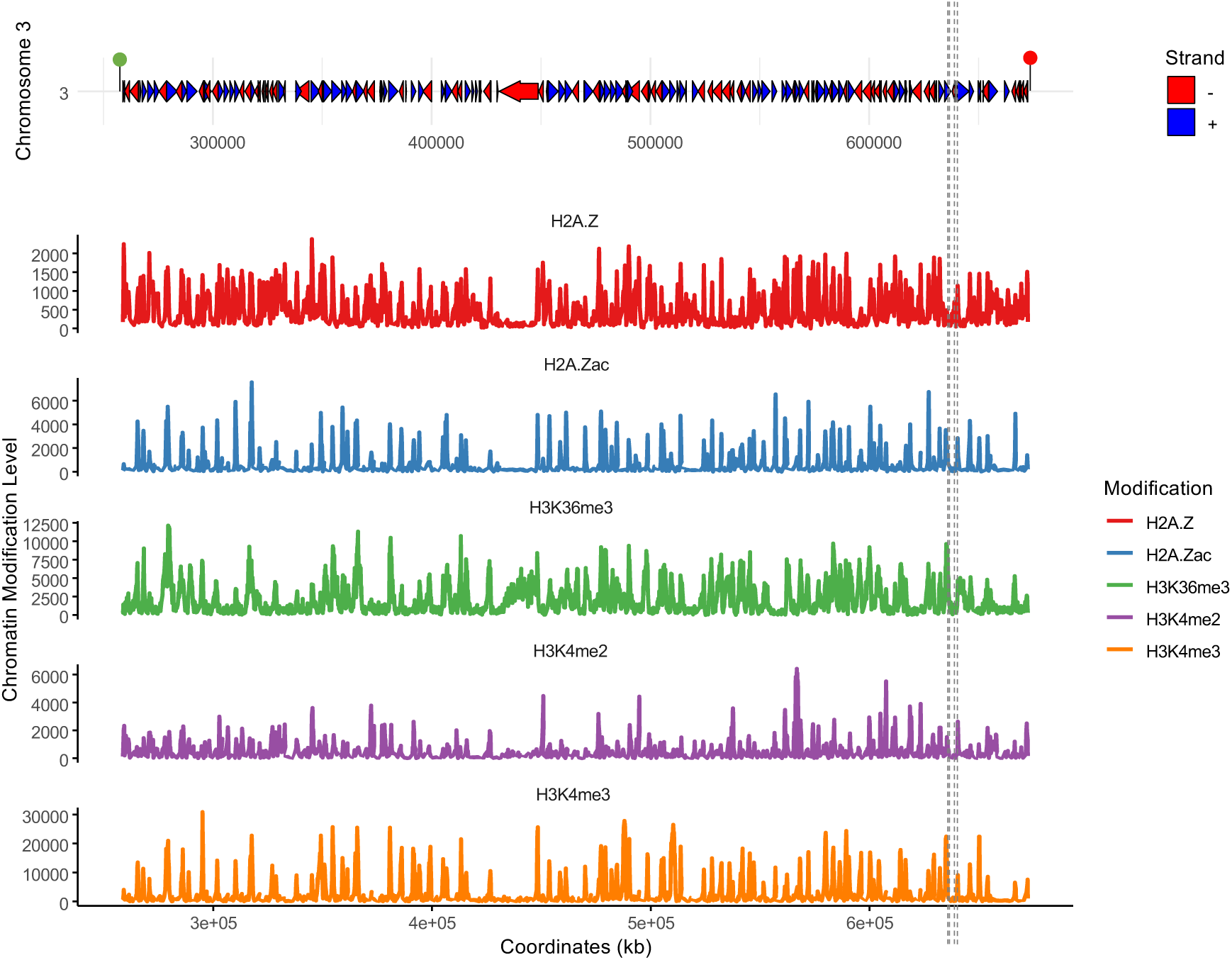
Gene structure and chromatin landscape within the *End3a* interval. Gene locations and orientations are represented by horizontal arrows (top panel). Vertical dashed lines indicate the approximate positions of the gRNAs used. Levels of specific chromatin modifications are displayed in 100 bp windows. H2A.Z and H2A.Zac levels from Bieluszewski et al. (2022), H3K36me3 from Liu et al. (2021), H3K4me2 from Yu et al. (2021), H3K4me3 from Potok et al. (2022).

**Appendix Fig. S8.**
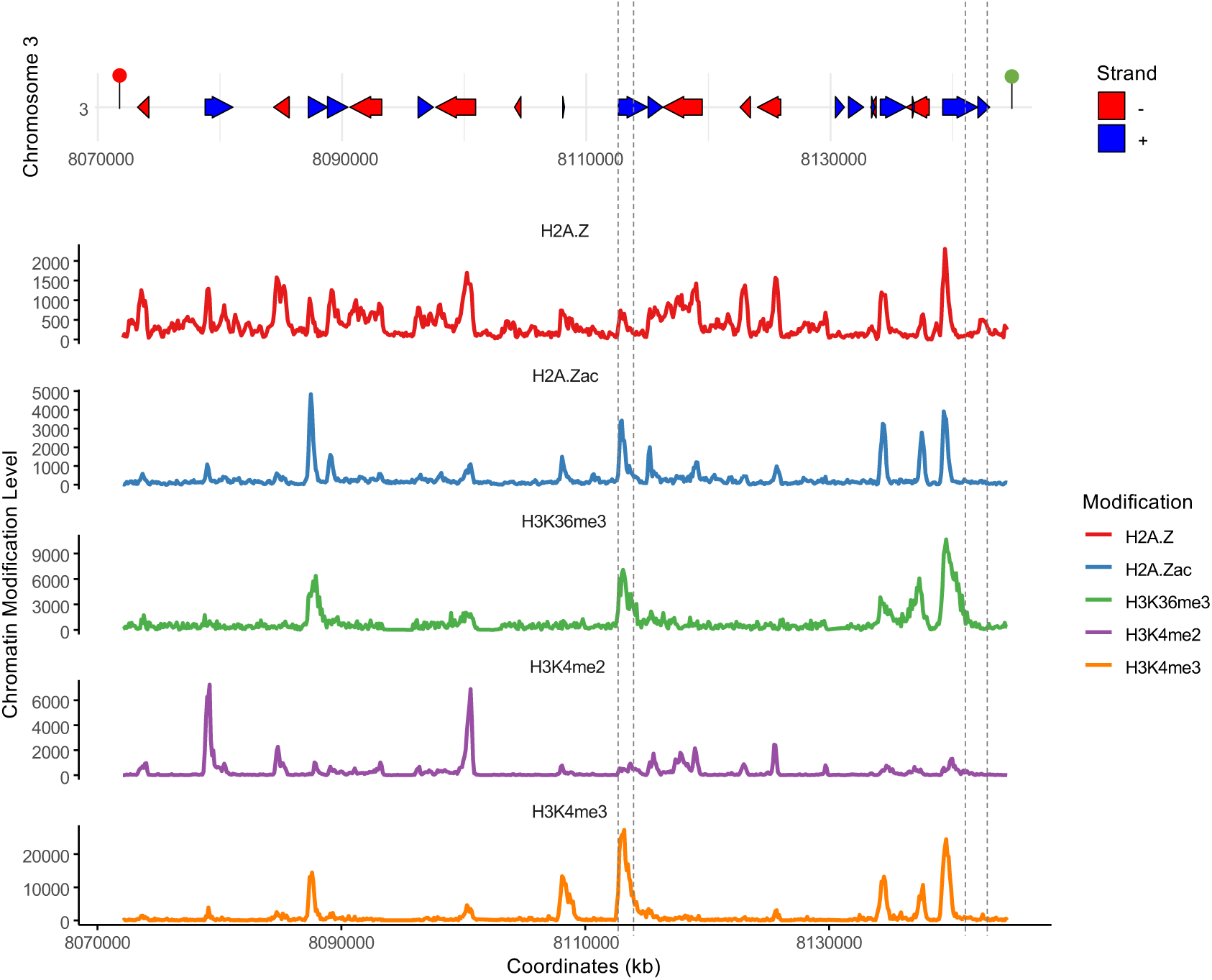
Gene structure and chromatin landscape within the *CW* interval. Gene locations and orientations are represented by horizontal arrows (top panel). Vertical dashed lines indicate the approximate positions of the gRNAs used. Levels of specific chromatin modifications are displayed in 100 bp windows. H2A.Z and H2A.Zac levels from Bieluszewski et al. (2022), H3K36me3 from Liu et al. (2021), H3K4me2 from Yu et al. (2021), H3K4me3 from Potok et al. (2022).

**Appendix Fig. S9.**
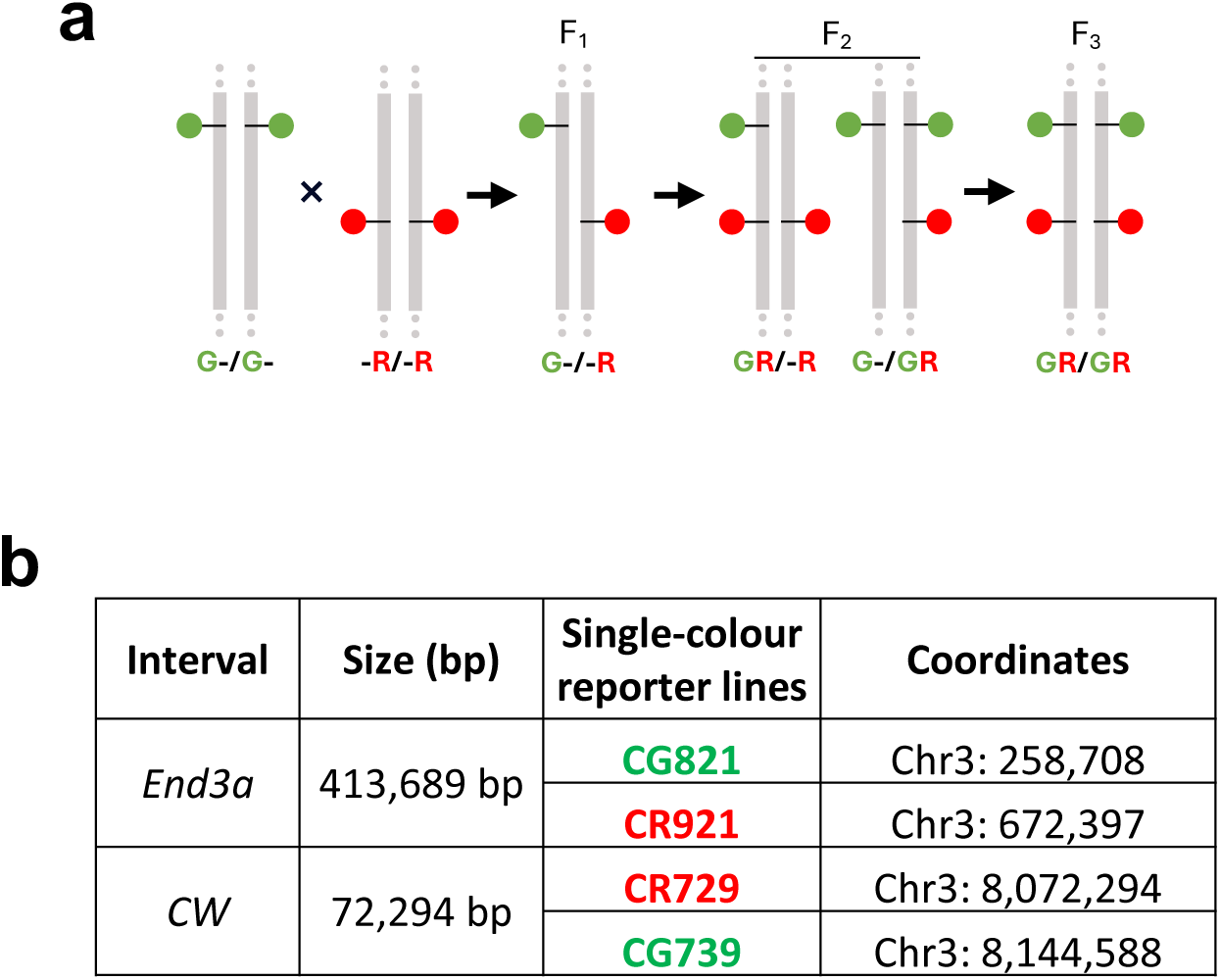
Generation of *End3a* and *CW* lines. **a**, Schematic overview of line generation: Plants carrying single red or green reporters were crossed, and F_2_ progeny obtained by self-pollination of the F_1_ generation were screened for individuals fixed for one reporter and heterozygous for the other. In the subsequent generation, plants homozygous for both reporters were selected, forming the final *End3a* and *CW* lines. **b**, Table summarizing the key characteristics of the *End3a* and *CW* lines.

**Appendix Fig. S10.**
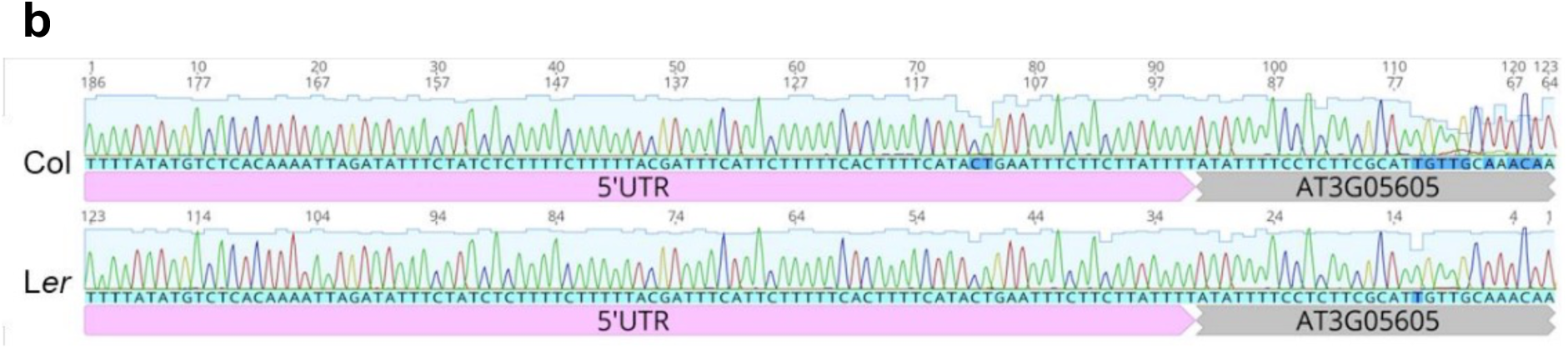
Identification of the Transcription Start Site (TSS) of the *CocoRNA* gene using 5’ RACE. 5’ UTR sequences of *CocoRNA* (AT3G05605) in Col (top) and L*er* (bottom).

**Appendix Fig. S11.**
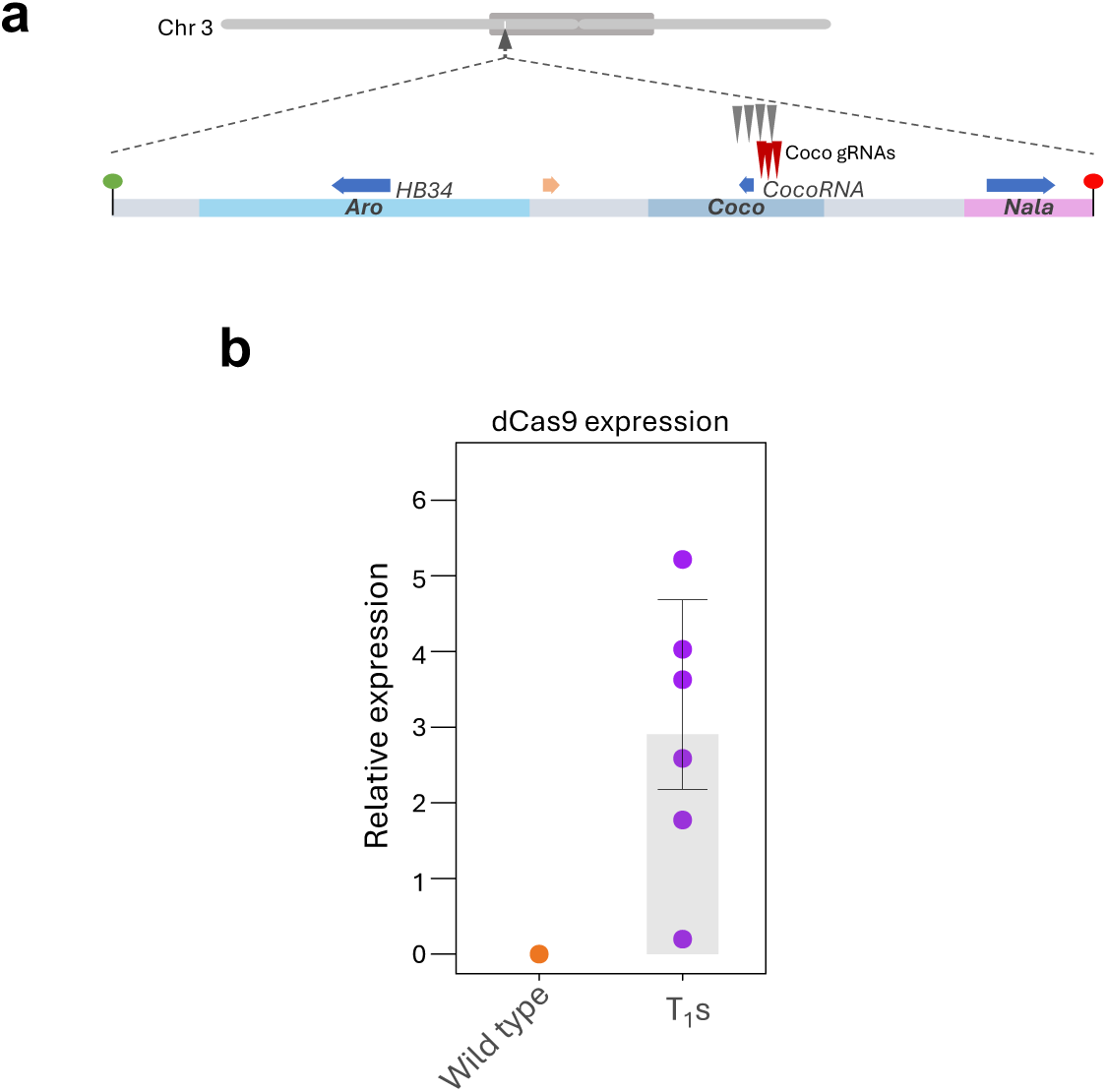
Targeting VP64 to the 5’ UTR of *CocoRNA*. a. Localization of gRNAs used for targeting dCas9-VP64 within the *ChP* interval. The top panel shows the position of *ChP* on the chromosome, with the pericentromeric region indicated by a dark gray oval. A zoomed-in view highlights the interval, marking the *Aro*, *Coco*, and *Nala* crossover hotspots. Blue horizontal arrows indicate gene locations, while an orange arrow marks a pseudogene. Maroon arrowheads represent gRNAs used for VP64 targeting, while gray arrowheads indicate gRNAs used for JMJ14 targeting (for comparison). b. Expression levels of dCas9-VP64 in T_1_ plants determined by RT-qPCR, normalized to *ACT2*. Each data point represents an independent T_1_ plant measurement.

